# StarBeast3: Adaptive Parallelised Bayesian Inference of the Multispecies Coalescent

**DOI:** 10.1101/2021.10.06.463424

**Authors:** Jordan Douglas, Cinthy L. Jiménez-Silva, Remco Bouckaert

## Abstract

As genomic sequence data becomes increasingly available, inferring the phylogeny of the species as that of concatenated genomic data can be enticing. However, this approach makes for a biased estimator of branch lengths and substitution rates and an inconsistent estimator of tree topology. Bayesian multispecies coalescent methods address these issues. This is achieved by embedding a set of gene trees within a species tree and jointly inferring both under a Bayesian framework. However, this approach comes at the cost of increased computational demand. Here, we introduce StarBeast3 – a software package for efficient Bayesian inference of the multispecies coalescent model via Markov chain Monte Carlo. We gain efficiency by introducing cutting-edge proposal kernels and adaptive operators, and StarBeast3 is particularly efficient when a relaxed clock model is applied. Furthermore, gene tree inference is parallelised, allowing the software to scale with the size of the problem. We validated our software and benchmarked its performance using three real and two synthetic datasets. Our results indicate that StarBeast3 is up to one-and-a-half orders of magnitude faster than StarBeast2, and therefore more than two orders faster than *BEAST, depending on the dataset and on the parameter, and is suitable for multispecies coalescent inference on large datasets (100+ genes). StarBeast3 is open-source and is easy to set up with a friendly graphical user interface.

## 1 Introduction

Existing methods for testing macro-evolutionary and macro-ecological questions have not kept pace with the explosion of next generation sequence data now available (Blom et al., 2016b; Bragg et al., 2017; Stenson et al., 2017). Despite burgeoning databases of within- and between-species genomic diversity (Blom et al., 2016b; Bragg et al., 2017; Stenson et al., 2017), it is still common practice to ignore the gene tree discordance that underlies any species phylogeny inferred from multi-locus sequences and instead infer species ancestry based on concatenated sequence data taken to represent all underlying gene histories (Degnan and Rosenberg, 2009; Heled and Drummond, 2010; Ogilvie et al., 2017; Jones, 2017; Rannala and Yang, 2017). While this approach can perform well when branches are long and incomplete lineage sorting (ILS) is absent, these conditions are rarely met.

Species trees inferred from concatenated sequences are often topologically incorrect (Degnan and Rosenberg, 2009; Heled and Drummond, 2010; Ogilvie et al., 2017), provide biased estimates for branch lengths and substitution rates (Ogilvie et al., 2016; Mendes and Hahn, 2016), and underestimate uncertainty in tree topology, resulting in an unjustified degree of confidence in the wrong tree (Heled and Drummond, 2010; Ogilvie et al., 2017). Such biases are exacerbated by subsampling of incongruent genes (Edwards et al., 2016; Mendes and Hahn, 2016), and hold even for deep splits in the tree (Oliver, 2013). These are crucial concerns in themselves and, more generally, can lead to biased estimates and erroneous inferences about fundamental evolutionary and ecological processes that require accurate phylogenetic trees, such as rates of speciation and extinction (Rowe et al., 2011; Cadena et al., 2011; Pepper et al., 2013), rates of substitution in DNA sequences (Bouckaert et al., 2013) and morphological characters (Pepper et al., 2013), species ancestry and ancestral age estimation (Mitchell et al., 2014), geographical history and origins (Lemey et al., 2009; Bouckaert, 2016), and species delimitation (Grummer et al., 2013; Yang and Rannala, 2014; Leaché et al., 2014; Yang and Rannala, 2010).

The multispecies coalescent (MSC; Maddison (1997); Edwards (2009); Liu et al. (2009)) is an approach designed to minimise these potential biases by modelling macro-evolution as a distribution of gene trees embedded within a species tree (Degnan and Rosenberg, 2009; Heled and Drummond, 2010; Ogilvie et al., 2017; Jones, 2017; Rannala and Yang, 2017). In doing so, the MSC provides a more biologically realistic framework for phylogenetic inference that captures the process of ILS underlying most multi-locus phylogenies. Furthermore, by explicitly modelling both species and gene trees, the MSC can address questions that cannot be addressed under a concatenation approach – such as automatic species delimitation (Fujita et al., 2012), with important implications for biodiversity assessment and conservation (Bickford et al., 2007).

A number of software packages have implemented the MSC in various ways (see review by Liu et al. (2015)). Our work at the Centre for Computational Evolution at the University of Auckland has led the development of *BEAST (STARBeast; Heled and Drummond (2010)) and StarBeast2 (Ogilvie et al., 2017) – full Bayesian MSC frameworks for species tree estimation from multilocus sequence data – and UglyTrees for visualising these models (Douglas, 2020). By explicitly modelling the MSC and avoiding the biases associated with concatenation methods (Heled and Drummond, 2010; Ogilvie et al., 2017, 2016), an analysis using either of these software packages can significantly change the conclusions drawn from data.

However, despite some advances in computational efficiency of the full Bayesian MSC (Ogilvie et al., 2017; Jones, 2017; Rannala and Yang, 2017), these complex models remain computationally intractable for large next generation sequence datasets of 100’s of sequenced loci across hundreds of individuals (i.e., 10^4^–10^6^ samples×loci). As a result, existing applications of the approach have tended to consider smaller datasets (Kang et al., 2014; Blom et al., 2016a) or to ignore much of the available data (Blom et al., 2016b; Bragg et al., 2017; Stenson et al., 2017), which reduces accuracy and increases uncertainty in species tree estimates (Ogilvie et al., 2017; Song et al., 2012). One approach to this problem has been the development of much simpler summary coalescent methods which utilise distributions of estimated gene tree topologies as input to rapidly process large datasets (Liu et al., 2015). These include the rooted triplet method MP-EST (Liu et al., 2010) and the quartet method ASTRAL (Mirarab et al., 2014). However, summary coalescent methods are sensitive to gene tree errors (Mirarab and Warnow, 2015; Xi et al., 2015) and produce trees in coalescent units, and thus time and population size estimates used by downstream analyses are confounded.

**Fig. S1:**
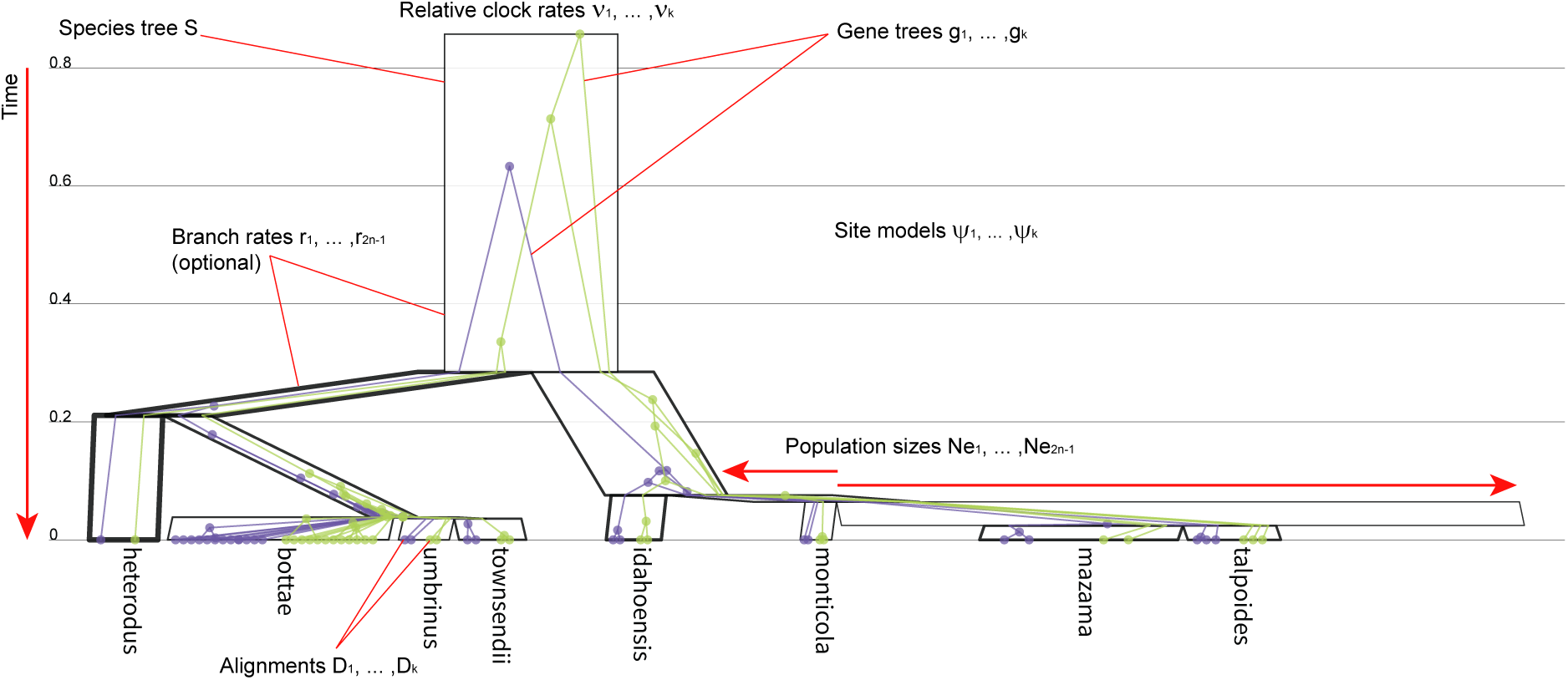
A species tree *S* with *n* = 8 species with *k* = 2 gene trees embedded. In this depiction, node heights (time) are shown on the y-axis and effective population sizes on the x-axis. Species tree branch rates are indicated by line thickness. Tree was built from a Gopher dataset (Belfiore et al., 2008) and visualised using UglyTrees (Douglas, 2020).

Here, we aim to perform Bayesian inference on large datasets using the Markov chain Monte Carlo (MCMC) algorithm as our workhorse. As illustrated in Fig. S1, the number of parameters involved is quite large, as is the accompanying state space. We develop a set of new MCMC proposals to explore state space in a much more efficient way than previous implementations and demonstrate we can handle datasets several times faster than *BEAST and StarBeast2. The resulting software package StarBeast3 is available as an open-source BEAST 2 package (Bouckaert et al., 2019).

## 2 Methods

### 2.1 The Multispecies Coalescent

Our objective is to develop efficient methods in a Bayesian framework for analysing models where there is a phylogeny, *S*, such as a species or language tree, that forms a constraint on a set of *k* trees **G** = {*g*_1_, …, *g*_*k*_}, such as gene trees. Each taxon within **G** is assigned to a single taxon within *S*, from some fixed individual-to-species mapping function (Fig.S1). Species tree *S* = (*T*_*S*_, *t*_*S*_) consists of a topology *T*_*S*_ and divergence times *t*_*S*_, as does the set of gene trees **G** = (*T*_**G**_, *t*_**G**_).

All trees are assumed to be binary rooted time trees, with fixed taxon node heights (typically extant; height 0). Each gene tree *g*_*i*_ consists of 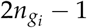 nodes and 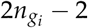 branches for taxon count 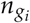, while *S* consists of 2*n*_*S*_ − 1 nodes and 2*n*_*S*_ − 1 branches, including a root branch, for species count *n*_*S*_. Gene tree taxa are associated with data *D* = {*D*_1_, …, *D*_*k*_}, e.g., nucleotide sequences or cognate data. Let *θ* be a set of model parameters, for instance those related to the speciation or nucleotide substitution processes. Consider the posterior density function *p*(*S*, **G**, *θ*|*D*):

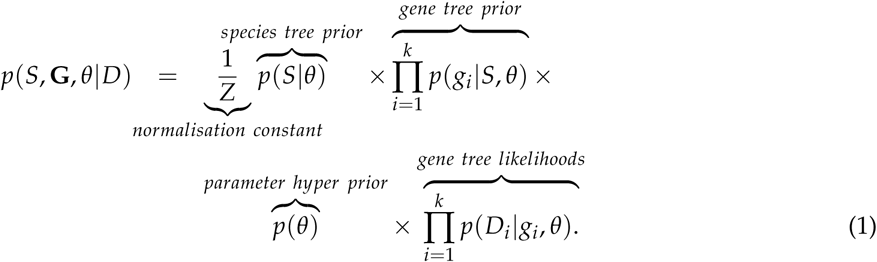

The MSC model is therefore hierarchical. *S* can follow a range of tree prior distributions *p*(*S*|*θ*), such as the Yule (Yule, 1925) or birth-death models (Nee et al., 1994). Whereas, each gene tree *g*_*i*_ is assumed to follow the multispecies coalescent process (Degnan and Rosenberg, 2009; Heled and Drummond, 2010; Ogilvie et al., 2017; Jones, 2017; Rannala and Yang, 2017), under which each species tree branch is associated with an independently and identically distributed (effective) population size **N**_**e**_ that governs the coalescent process of **G**, where |**N**_**e**_| = 2*n*_*S*_ −1. Gene trees are thus assumed to be contained within *S* (Fig. S1).

Site evolution is assumed to follow a continuous-time Markov process (Felsenstein, 1981) under some substitution model and clock model:

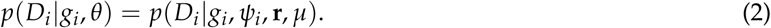

*ψ*_*i*_ can adopt a range of molecular substitution models, such as the HKY nucleotide evolution model (Hasegawa et al., 1985) or the WAG amino acid evolution model (Whelan and Goldman, 2001). Tree *g*_*i*_ has relative molecular substitution rate *ν*_*i*_ ∈ *ψ*_*i*_. Each branch in *S* is associated with a substitution rate **r** which governs the rate of site evolution of **G** along the respective branch, where |**r**| = 2*n*_*S*_ −1 (Fig. S1). Branch rates **r** are assumed to be independently and identically distributed under a log-normal distribution with standard deviation *σ* (i.e., the multispecies coalescent relaxed clock model (Ogilvie et al., 2017; Drummond et al., 2006)). Lastly, the clock rate *µ* can be estimated when accompanied by time-calibration data, such as ancient fossil records (Heled and Drummond, 2012; Sauquet et al., 2011; Ballesteros and Sharma, 2019), or left fixed when no such data is available. Overall, the total substitution rate of any given branch in *g*_*i*_ is the product of *ν*_*i*_, *µ*, and a subset of the elements in **r** (weighted by their coverage of the gene tree branch; Ogilvie et al. (2017)).

In this article, we develop tools that allow the MSC to be applied to large datasets using complex models of evolution. Although we focus on MSC models, we anticipate that in the future other models of the form expressed in Eq. (1) will be developed, e.g., models that allow some lateral gene transfer and therefore allow some gene tree branches to cross species boundaries in the species tree. We design a number of MCMC operators which generate proposals that explore the state space more efficiently – using a Gibbs sampler for population sizes, a combination of Bactrian (Yang and Rodríguez, 2013; Thawornwattana et al., 2018) and adaptable variance multivariate normal (Baele et al., 2017) proposal kernels, a parallel operator for sampling gene trees and substitution model parameters, and an MCMC operator which selects other operators based on their exploration efficiency (Douglas et al., 2021b). Moreover, in the special case of the multispecies relaxed clock model (Ogilvie et al., 2017), we introduce methods for operating on the species tree, the gene trees, and the clock model simultaneously (Zhang and Drummond, 2020; Douglas et al., 2021b).

### 2.2 Effective Population Size Gibbs Operator

The StarBeast2 (Ogilvie et al., 2017) and DISSECT (Jones et al., 2015) packages have the capability of integrating out effective population sizes **N**_**e**_ when using an inverse gamma distributed prior on **N**_**e**_, based on a technique introduced by Liu et al. (2008) and detailed out by Jones (2017). This approach greatly reduces the state space. However, consequently the posterior Eq. 1 can no longer be broken down in a product over components over individual gene trees:

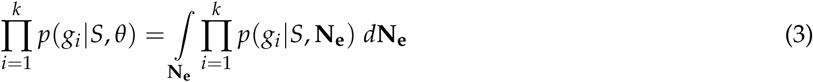

Thus, the technique is not suitable for gene-tree operator parallelisation, and therefore we estimate **N**_**e**_ instead.

Suppose that 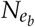, for species-tree branch *b*, follows an inverse gamma prior distribution Inv-Γ(*α*_*N*_, *µ*_*N*_), where the shape *α*_*N*_ is fixed at 2 and therefore the scale *µ*_*N*_ is the expected value (because 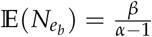). Following the results by Jones (2017), the posterior of 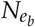 follows an inverse gamma Inv-Γ(*α*_*N*_*′, µ*_*N*_*′*), such that *α*_*N*_*′* = *α*_*N*_ + *a* and *µ*_*N*_*′* = *µ*_*N*_ + *c* where *a* is the total number of coalescent events of all gene trees in branch *b* and 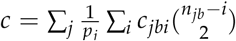. Here *p*_*j*_ is the ploidy of gene *g*_*j*_, *c*_*jbi*_ the size of the *i*th coalescent interval for gene *g*_*j*_ in branch *b*, and *n*_*jb*_ the number of lineages of gene tree *g*_*j*_ at the tip-side of branch *b* (so that *n*_*jb*_ −*i* is the number of lineages at the start of the *i*th coalescent interval for *g*_*j*_).

Instead of integrating out **N**_**e**_, our `GibbsPopulation` operator samples from the posterior. All 2*n*_*S*_ −1 elements in **N**_**e**_ are proposed simultaneously. As demonstrated later, this turns out to be more efficient than standard **N**_**e**_ random walk operators, with the added advantage of sampling effective population sizes – which may be a parameter of interest – as well as the ability to parallelise gene tree proposals. This technique is readily applicable for periodically sampling and logging **N**_**e**_ to implementations that do integrate this term out.

### 2.3 Bactrian Operators for Trees

The step size of a proposal kernel should be such that the proposed state *x′* is sufficiently far from the current state *x* to explore vast areas of parameter space, but not so far that the proposal is rejected too often (Roberts et al., 1997). The Bactrian distribution (Yang and Rodríguez, 2013; Thawornwattana et al., 2018) has minimal probability mass around the center, and a higher concentration flanking the center, akin to the humps of a Bactrian camel (Fig. S2; left). This distribution is a preferred alternative to standard uniform- or normal-distributed random walk kernels, as it places minimal probability on step sizes that are too large or too small, and has successfully improved phylogenetic inference in previous studies (Yang and Rodríguez, 2013; Zhang and Drummond, 2020; Douglas et al., 2021b).

**Fig. S2:**
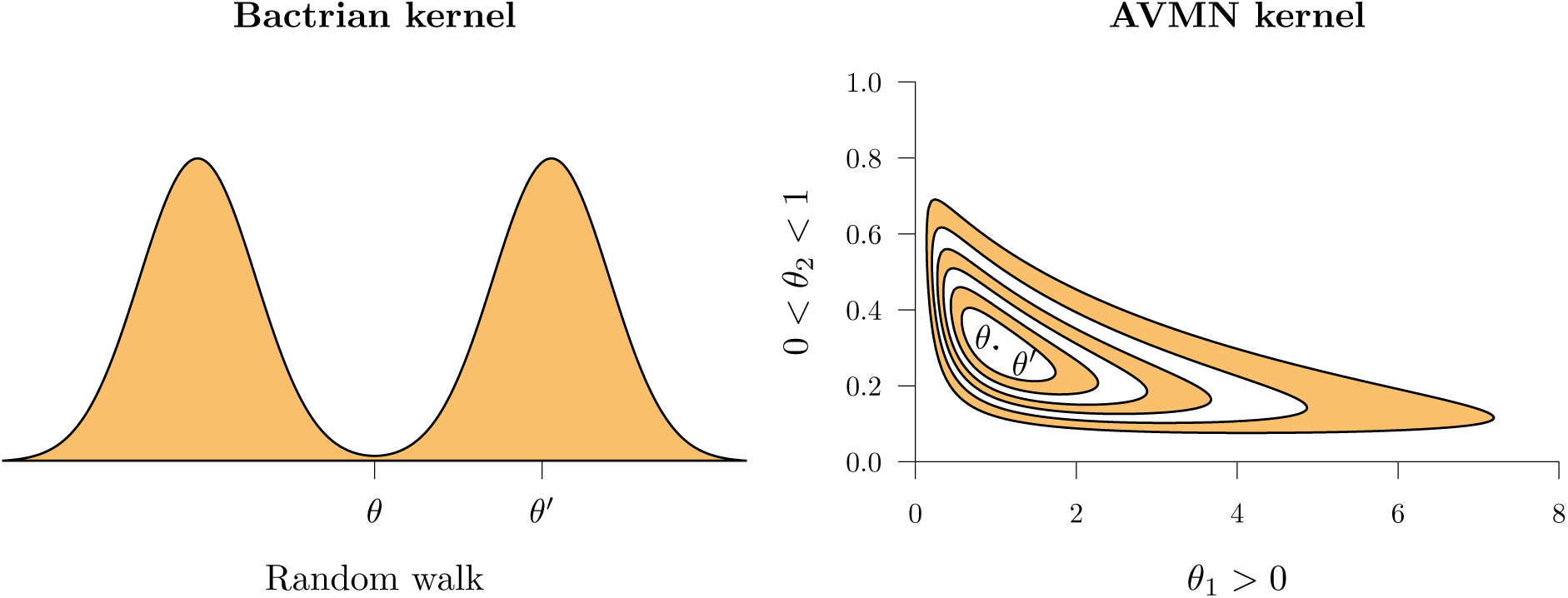
Depiction of random walks *θ* → *θ′* under varying proposal kernels. Left: The random walk occurs from the origin between the two modes, where the vertical axis shows the probability density function of the kernel *p*(*θ′*|*θ*) (Yang and Rodríguez, 2013). Right: A two dimensional random walk on inversely correlated parameters *θ* = (*θ*_1_, *θ*_2_) with different domains (Baele et al., 2017). Contours describe the joint probability density function *p*(*θ*_1_, *θ*_2_) under a transformed multivariate normal distribution learned during MCMC.

In this article, we apply Bactrian proposals to trees. The standard set of tree node height proposals in BEAST 2 consists of a `Scale` operator which embarks all nodes in the tree on a random walk (in log-space), a `RootScale` operator which does so for only the root of a tree, an `UpDown` operator which changes species/gene node heights and various continuous parameters simultaneously (Drummond et al., 2002) a `SubtreeSlide` operator which slides a node up or down branches (Hohna et al., 2008), and constant distance operators when a relaxed clock model is applied (Zhang and Drummond, 2020). Each operator would normally draw a random variable from a uniform distribution, but here we instead use a Bactrian distribution and apply appropriate transformations. We also introduce the `Interval` operator, which transforms parameters with lower- and upper-bounds (such as tree node heights) by applying a Bactrian random walk in their real-space transformations.

### 2.4 AVMN Operator

An adaptive variance multivariate normal (AVMN) operator (Baele et al., 2017) provides proposals for a set of real-space parameters by learning the posterior throughout the run of the MCMC algorithm and approximating it as a multivariate normal distribution to capture correlations between parameters (Fig. S2; right). The space spanned by such a set continuous parameters may need to be transformed to satisfy the real-space assumption, by applying a log-transformation to parameters with positive domains (such as substitution rates), or a log-constrained sum transformation to multivariate parameters with unit sums (such as nucleotide frequencies), for instance. AVMN has been demonstrated to be more efficient in estimating phylogenetic parameters than standard random walk or scale operators (Baele et al., 2017; Bouckaert, 2020; Douglas et al., 2021b).

Consider a single gene tree *g*_*i*_ and its substitution model *ψ*_*i*_, consisting of substitution rates and nucleotide frequencies for instance. Performing a single proposal for any single parameter would require a full recalculation of the tree likelihood *p*(*D*_*i*_|*g*_*i*_, *ψ*_*i*_, **r**, *µ*) (see peeling algorithm by Felsenstein (1981)). Therefore, proposing all site model parameters *ψ*_*i*_ simultaneously can reduce the number of likelihood calculations required and thus lower the computational runtime.

### 2.5 Parallel Gene Tree Operator

During MCMC, operators are typically sampled proportionally to fixed weights (or proposal probabilities), to ensure the chain is ergodic. Here, we present an alternative method, where a single gene tree *g*_*i*_ and its substitution model *ψ*_*i*_ is selected, and *N*_*p*_ operators are sequentially sampled and applied to *g*_*i*_ and *ψ*_*i*_, before returning to the full parameter space. This is equivalent to running a small MCMC chain of *N*_*p*_ steps – applying only gene tree and substitution model operators on *g*_*i*_ and *ψ*_*i*_ – and then accepting the resulting *g*_*i*_*′* and *ψ*_*i*_*′* afterwards with probability 1, as if it were a single Gibbs sampling operation (Geman and Geman, 1984).

Observe that because only *g*_*i*_ and its associated parameters change, part of Eq (1) can be rewritten as:

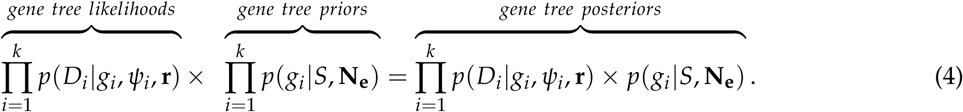

Thus, the posterior distribution can be decomposed into the product of contributions of individual gene trees and their substitution models. Assuming that substitution model parameters *ψ*_*i*_ are distinct for each gene tree *g*_*i*_, an *N*_*p*_-step MCMC chain could be run for each of *g*_*i*_ and *g*_*j*_ for (*i* ≠ *j*) in parallel, and the resulting *g*_*i*_*′* and *g*_*j*_*′* each accepted with probability 1, as if two Gibbs operators were sequentially applied. Because the posterior density for *g*_*i*_ is proportional to *p*(*D*_*i*_|*g*_*i*_, *θ*) *p*(*g*_*i*_|*S, θ*) and that of *g*_*j*_ proportional to *p*(*D*_*j*_ |*g*_*j*_, *θ*) *p*(*g*_*j*_ |*S, θ*), then provided that any shared parameters (such as **r**, *S*, and **N**_**e**_) are not being operated on, these two *N*_*p*_-step MCMC chains can run in parallel.

Where there are *N*_*t*_ threads available, the *k* gene trees are split into *N*_*t*_ groups (assuming *k* ≥ *N*_*t*_). The *N*_*t*_ sets of *N*_*p*_-step MCMC chains are run in parallel and the resulting gene trees **g** are accepted into the main MCMC chain. Here, we introduce a parallel operator `ParallelGeneTreeOperator(`**G**, ◼). This operator partitions gene trees into *N*_*t*_ threads and operates on their topologies, node heights, and substitution models. Tree node height proposals employ the Bactrian kernel where applicable (Fig. S2), and substitution model proposals invoke the AVMN kernel (Fig. S2). The chain length *N*_*p*_ of each thread is learned during MCMC (Fig. S3).

Since each small MCMC chain for a thread can be considered a single Gibbs proposal, for *N*_*t*_ threads in principle *N*_*t*_ steps should be added to the main chain. If the operator is selected just before logging a state, in principle some threads may need to be disregarded before logging in order to maintain exactly equal intervals in the trace log. Due to the low frequency at which the operator is selected, and the logging intervals being orders of magnitude larger than the number of threads, this does not appear to be a problem in practice.

### 2.6 Species Tree Relaxed Clock Model Operators

The constant distance operator family exploits the negative correlations between divergence times and branch substitution rates by proposing both terms simultaneously (Zhang and Drummond, 2020). This technique has yielded a parameter convergence rate one to two orders of magnitude faster, particularly for large datasets that come with peaked posterior distributions (Douglas et al., 2021b). Under the multispecies coalescent relaxed clock model used by StarBeast2, the branch rate of gene tree branch *b* is the length-weighted branch rate **r** of all species tree branches that contain *b* (Ogilvie et al., 2017). Moreover, effective population sizes **N**_**e**_ are positively correlated with divergence times, so this correlation could also be readily exploited.

Extending the work by Zhang and Drummond (2020), we introduce the `ConstantDistanceMSC` operator. This operator proposes a node height *t*_*X*_ for species tree internal node *X*, the three branch rates **r** and population sizes **N**_**e**_ incident to *X*, and heights for all gene tree non-leaf nodes that are contained within these three incident branches (Fig. S4). *t*_*X*_ is embarked on a Bactrian random walk (Yang and Rodríguez, 2013) to give *t′*_*X*_, then **r** and the node heights in **G** are proposed such that all genetic distances are conserved following the change in *t*_*X*_, and **N**_**e**_ is proposed such that the positive correlation between itself and the branch lengths incident to *X* is respected (see Algorithm S1).

Previously, we introduced the narrow exchange rate (`NER`) operator (Douglas et al., 2021b). This operator combined the simple `NarrowExchange` operator (i.e., a proposal which swaps a subtree with its uncle subtree; Drummond et al. (2002)) with the `ConstantDistance` operator (Zhang and Drummond, 2020), by applying a small topological change to the tree and then recomputing branch substitution rates such that evolutionary distances are preserved. We demonstrated that this operator assisted the traversal of tree topology space on longer alignments compared with shorter ones.

**Fig. S3:**
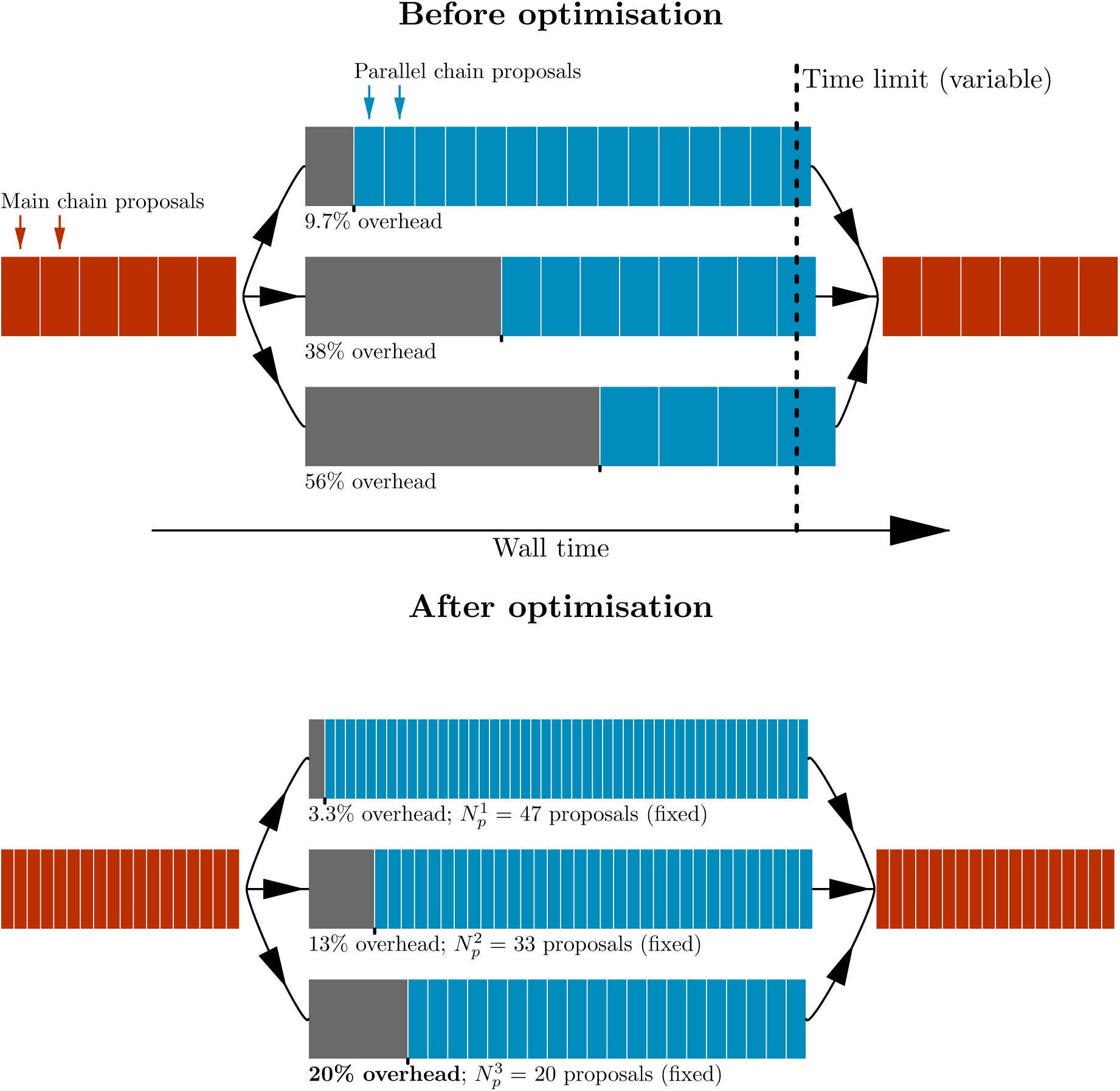
Optimisation of gene tree parallel operator chain lengths. Top: The time limit of each parallel MCMC chain is randomised on each call so that the overhead (intercept) and time-per-proposal (slope) can be learned as a linear regression model. Bottom: The linear regression model is applied, and parallel MCMC chain lengths are set such that the slowest thread attains the user-specified target overhead (i.e., the bottom thread has attained 20% overhead in the example above).

**Fig. S4:**
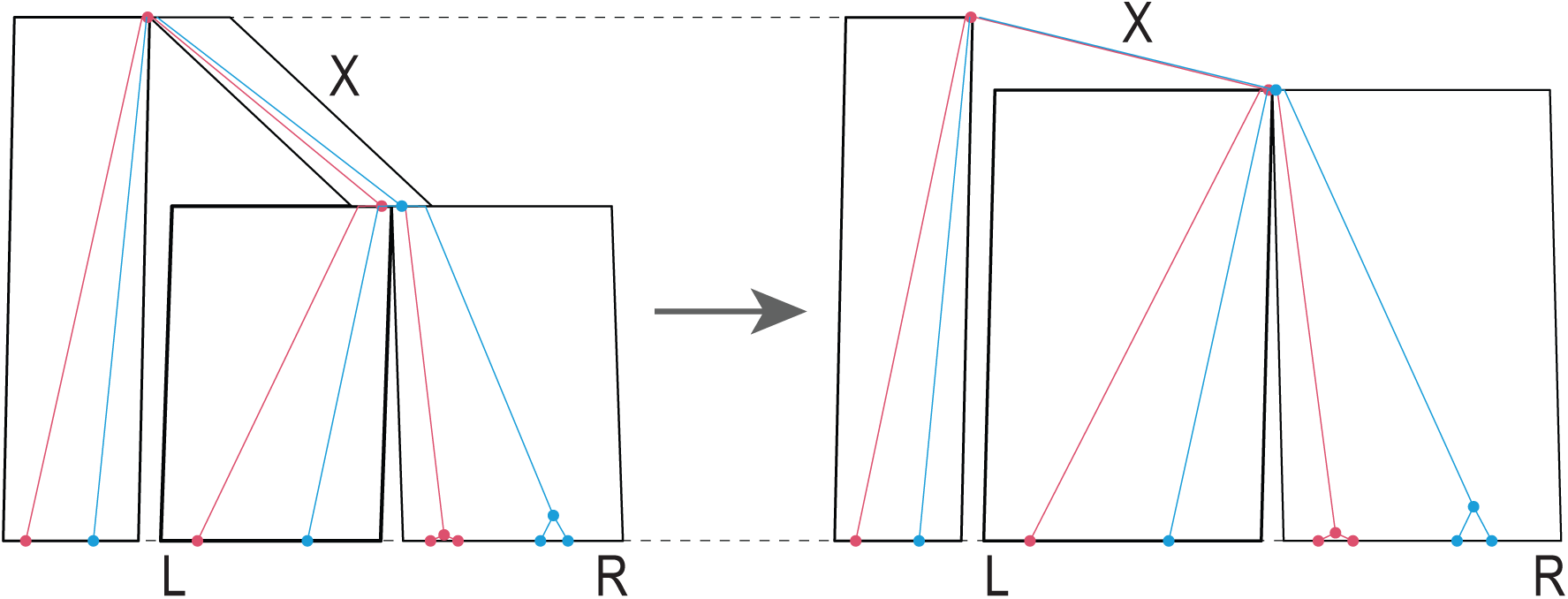
An example of an `ConstantDistanceMSC` proposal, with species nodes *X*, and its two children *L* and *R* indicated. The height of *X* is increased, and **r** (indicated by species node line thickness) and **N**_**e**_ (indicated by species node width) are increased or decreased accordingly. Figure generated by UglyTrees (Douglas, 2020).

Here, we combine this work with the `CoordinatedExchange` operator implemented by Ogilvie et al. (2017) – based on work by Jones (2017) and Rannala and Yang (2017) – and introduce the coordinated narrow exchange rate (`CNER`) operator. This operator exchanges a species tree node with its uncle node, adjusts gene tree topologies **g** to preserve compatibility with *S*, and proposes three nearby branch rates **r** to preserve genetic distances (Algorithm S2).

### 2.7 Adaptive Operator Weighing

Previously, we developed the `AdaptableOperatorSampler(`*x*) operator (Douglas et al., 2021b). This operator learns the weights (or proposal probabilities) behind a set of sub-operators during MCMC, by rewarding operators which bring about large changes to parameter *x* in short computational runtime, with respect to some distance function: Euclidean distance when *x* is real, and RNNI distance (Collienne and Gavryushkin, 2021) when *x* is tree topology. This approach can account for the scenario when an operator’s performance is conditional on the dataset. When a dataset contains very little signal with respect to a certain parameter *x* and its prior distribution, then resampling that parameter from its prior distribution using the `SampleFromPrior (`*x*) operator may be more efficient than embarking *x* on a random walk, for instance (Douglas et al., 2021b). In contrast, datasets with more signal are likely to prefer smarter operators which account for correlations in the posterior distribution, such as the constant distance or `NER` operators (Zhang and Drummond, 2020; Douglas et al., 2021b).

Here, we have applied the `AdaptableOperatorSampler` to seven areas of parameter space: the species and gene tree node heights (*t*_*s*_ and *t*_**G**_), the relaxed clock model rates **r** and standard deviation *σ*, the mean effective population size *µ*_*N*_, the species tree birth rate *λ* (assuming a Yule speciation model (Yule, 1925)), and the species tree topology *T*_*S*_. These operator schemes are explicated in **Tables S1** and **S2**.

**Table S1:** StarBeast3 operator scheme, assuming a Yule tree prior on the species tree with birth rate *λ* (Yule, 1925). The `ParallelMCMC` operator weight was set such that it consists of 1% of the all proposals. Further operator details can be found in Drummond and Bouckaert (2015). ^†^Bactrian kernel applied to random walk (Yang and Rodríguez, 2013).

| Operator | Weight | Reference |
| --- | --- | --- |
| **Species tree** |  |  |
| NodeReheight($T_S, t_S, T_G, t_G$) | 30 | Ogilvie et al. (2017) |
| CoordinatedUniform($t_S, t_G$) | 30 | Ogilvie et al. (2017); Jones (2017) |
| CoordinatedExponential($t_S, t_G$) | 15 | Ogilvie et al. (2017); Jones (2017) |
| SubtreeSlide($T_S, t_S$)<sup>†</sup> | 15 | Hohna et al. (2008) |
| WilsonBalding($T_S$) | 15 | Drummond et al. (2002) |
| WideExchange($T_S$) | 15 | Drummond et al. (2002) |
| AdaptableOperatorSampler($T_S$) | 15 | |
| NarrowExchange($T_S$) | | Drummond et al. (2002) |
| CoordinatedExchange($T_S, T_G$) | | Ogilvie et al. (2017) |
| NER($T_S, \mathbf{r}$) | | Douglas et al. (2021b) |
| CNER($T_S, \mathbf{r}, T_G$) | | *Species Tree Relaxed Clock Model Operators* |
| Uniform($t_S$)<sup>†</sup> | 3 | |
| RootScale($t_S$)<sup>†</sup> | 3 | |
| Interval($t_S$)<sup>†</sup> | 3 | *Bactrian Operators for Trees* |
| AdaptableOperatorSampler($t_S$) | 100 | |
| Uniform($t_S$)<sup>†</sup> | | |
| TreeScale($t_S$)<sup>†</sup> | | |
| Interval($t_S$)<sup>†</sup> | | *Bactrian Operators for Trees* |
| ConstantDistanceMSC($t_S, t_G, \mathbf{r}, \mathbf{N}_e$)<sup>†</sup> | | *Species Tree Relaxed Clock Model Operators* |
| CoordinatedUniform($t_S, t_G$) | | Ogilvie et al. (2017); Jones (2017) |
| CoordinatedExponential($t_S, t_G$) | | Ogilvie et al. (2017); Jones (2017) |
| UpDown($[t_S, t_G, \mathbf{N}_e, \mu_N], [\cdot, \lambda, \mu]$)<sup>†</sup> | | Drummond et al. (2002) |
| **Gene trees / site models** |  |  |
| ParallelMCMC($T_G, t_G, \psi$) | 3.42 | **Table S2** |
| **Tree hyperparameters** |  |  |
| GibbsPopulation($\mathbf{N}_e$) | 50 | *Effective Population Size Gibbs Operator* |
| AdaptableOperatorSampler($\mu_N$) | 5 | |
| Scale($\mu_N$)<sup>†</sup> | | |
| UpDown($[t_S, t_G, \mathbf{N}_e, \mu_N], [\cdot, \lambda, \mu]$)<sup>†</sup> | | Bouckaert et al. (2019) |
| SampleFromPrior($\mu_N$) | | Douglas et al. (2021b) |
| AdaptableOperatorSampler($\lambda$) | 5 | |
| Scale($\lambda$)<sup>†</sup> | | |
| UpDown($[t_S, t_G, \mathbf{N}_e, \mu_N], [\cdot, \lambda, \mu]$)<sup>†</sup> | | Bouckaert et al. (2019) |
| SampleFromPrior($\lambda$) | | Douglas et al. (2021b) |
| **Relaxed clock model** |  |  |
| AdaptableOperatorSampler($\mathbf{r}$) | 30 | |
| Scale($\mathbf{r}$)<sup>†</sup> | | |
| ConstantDistanceMSC($t_S, t_G, \mathbf{r}, \mathbf{N}_e$)<sup>†</sup> | | *Species Tree Relaxed Clock Model Operators* |
| SampleFromPrior($\mathbf{r}$) | | Douglas et al. (2021b) |
| AdaptableOperatorSampler($\sigma$) | 5 | |
| Scale($\sigma$)<sup>†</sup> | | |
| SampleFromPrior($\sigma$) | | Douglas et al. (2021b) |

**Table S2:** StarBeast3 parallel operator scheme for gene trees and their associated site models (assumed to be an HKY model with transition-transversion ratio *κ* and nucleotide frequencies *f*). Each operator is applicable to a single gene tree *g*_*i*_ or its site model *ψ*_*i*_. 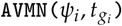 generated proposals for the site model and complete set of tree node heights simultaneously. Operator weights are normalised into proposal probabilities within a single MCMC chain called by `ParallelMCMC`. Further operator details can be found in Drummond and Bouckaert (2015). ^†^Bactrian kernel applied to random walk (Yang and Rodríguez, 2013).

| Operator | Weight | Reference |
| --- | --- | --- |
| $\text{ParallelMCMC}(T_G, t_G, \psi)$ | | *Parallel Gene Tree Operator* |
| **Gene trees** |  |  |
| $\forall_{i \in 1, \dots, k} \text{WilsonBalding}(T_{g_i})$ | 15 | Drummond et al. (2002) |
| $\forall_{i \in 1, \dots, k} \text{WideExchange}(T_{g_i})$ | 15 | Drummond et al. (2002) |
| $\forall_{i \in 1, \dots, k} \text{NarrowExchange}(T_{g_i})$ | 15 | Drummond et al. (2002) |
| $\forall_{i \in 1, \dots, k} \text{SubtreeSlide}(T_{g_i}, t_{g_i})^\dagger$ | 10 | Hohna et al. (2008) |
| $\forall_{i \in 1, \dots, k} \text{Uniform}(t_{g_i})$ | 30 | |
| $\forall_{i \in 1, \dots, k} \text{RootScale}(t_{g_i})^\dagger$ | 10 | |
| $\forall_{i \in 1, \dots, k} \text{Interval}(t_{g_i})^\dagger$ | 10 | *Bactrian Operators for Trees* |
| $\forall_{i \in 1, \dots, k} \text{AdaptableOperatorSampler}(t_{g_i})$ | 100 | |
| $\text{TreeScale}(t_{g_i})^\dagger$ | | |
| $\text{Uniform}(t_{g_i})^\dagger$ | | |
| $\text{SubtreeSlide}(T_{g_i}, t_{g_i})^\dagger$ | | Hohna et al. (2008) |
| $\text{EpochOperator}(t_{g_i})^\dagger$ | | Bouckaert (2021) |
| **Site models** |  |  |
| $\forall_{i \in 1, \dots, k} \text{AVMN}(\psi_i, t_{g_i})$ | 5 | Baele et al. (2017) |
| $\forall_{i \in 1, \dots, k} \text{Scale}(\kappa_i)^\dagger$ | 0.5 | |
| $\forall_{i \in 1, \dots, k} \text{Scale}(\nu_i)^\dagger$ | 0.5 | |
| $\forall_{i \in 1, \dots, k} \text{DeltaExchange}(f_i)$ | 0.5 | |

## 3 Results

In this section we prove StarBeast3 is correctly implemented through a well-calibrated simulation study. Further, we demonstrate that StarBeast3 is efficient at doing Bayesian inference on large datasets compared with StarBeast2. We did not compare to *BEAST directly, since it does not provide relaxed clock models on species trees, but note that Ogilvie et al. (2017) benchmarked StarBeast2 against *BEAST for strict clocks and found StarBeast2 to be an order faster than *BEAST, so any gain over StarBeast2 will be more so over *BEAST.

### 3.1 The Implementation Is Correct

In order to validate the correctness of StarBeast3, we performed two well-calibrated simulation studies. These were achieved by simulating nucleotide alignments (of two varying sizes) using parameters directly sampled from the prior distribution, and then recovering the posterior estimates of these parameters by doing Bayesian inference on the simulated alignments using StarBeast3. For each study, the 95%-coverage of each parameter was approximately 95% (meaning that the true parameter estimate was within the 95% highest posterior density interval approximately 95% of the time). Therefore, these experiments provide confidence in StarBeast3’s correctness, and are presented in Fig. S5 and Section 4 of *Supplementary Material*.

### 3.2 Performance Benchmarking

We evaluated the performance of StarBeast3 for its ability to achieve multispecies coalescent parameter convergence in a Bayesian framework, compared with that of StarBeast2. This was measured by computing effective sample sizes (ESS) generated per hour during MCMC across multiple replicates of three real and two simulated datasets (**Table S3**). The ESS of any parameter should be over 200 in order to estimate its posterior distribution (Tracer; Rambaut et al. (2018)). To allow both software packages to perform at their best, effective population sizes were integrated out by StarBeast2, but were estimated by StarBeast3. This section provides a general comparison of StarBeast3 and StarBeast2, however the performances of individual operators can be found in Sections 5 and 6 of *Supplementary Material*.

**Fig. S5:**
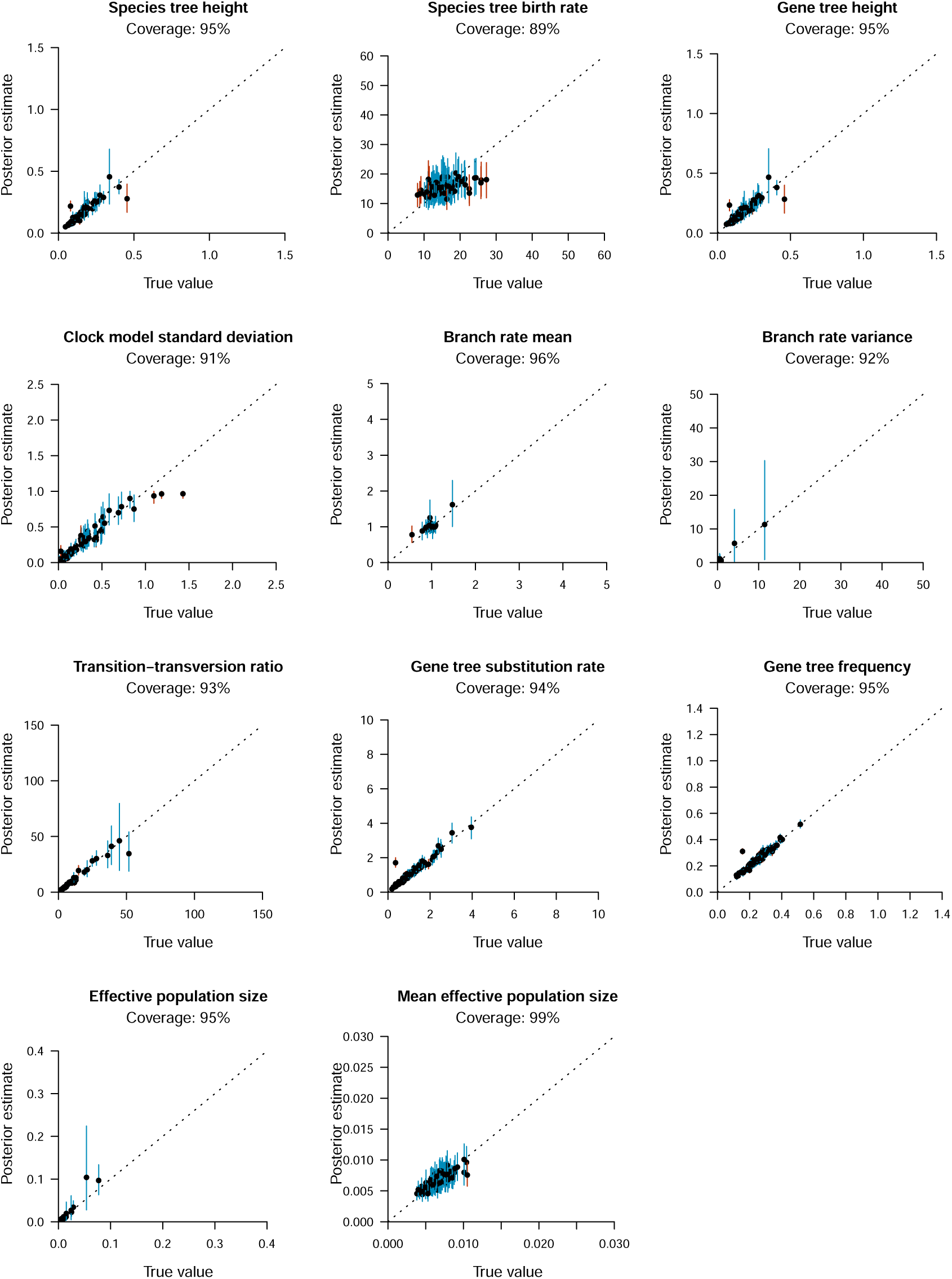
Well-calibrated simulation study analysing *n*_*S*_ = 16 species, *n*_**G**_ = 48 taxa, and *k* = 50 genes. 100 simulations were performed to recover the coverage between “true” simulated values and their estimates under the posterior distribution. 95% highest posterior density (HPD) intervals of parameters are represented by vertical lines. Each line represents a single simulation, and is coloured blue when the true value was contained within the 95% interval, or red otherwise. The top of each plot shows the coverage of each parameter (i.e., the number of MCMC simulations for which the “true” parameter value was contained within the 95% HPD).

**Table S3:** Benchmark datasets. 50 gene trees were subsampled from the Skink and Spider datasets. The Simulated datasets were directly sampled from the model specification used during Bayesian inference (described in Section 3 of *Supplementary Material*).

| Dataset | Number of species $n_S$ | Number of taxa $n_G$ | Number of gene trees $k$ |
| --- | --- | --- | --- |
| Frog (Barrow et al., 2014) | 21 | 88 | 26 |
| Skink (Bryson Jr et al., 2017) | 10 | 59 | 50 |
| Spider (Hamilton et al., 2016) | 36 | 83 | 50 |
| Simulated(12) | 4 | 12 | 100 |
| Simulated(48) | 16 | 48 | 100 |

The ESS/hr was evaluated in five distinct areas of parameter space. First, we considered generic summaries of convergence: the ESS/hr of the posterior density *p*(*θ* | *D*), the likelihood *p*(*D* | *θ*), and the prior density *p*(*θ*). Second, species tree *S* convergence was evaluated in terms of its height *h*_*S*_, its length *l*_*S*_, and hyperparameters *λ* – the Yule model birth rate (Yule, 1925) – and *µ*_*N*_ – the mean effective population size. In the case of StarBeast3, where effective population sizes are estimated, we also measured the mean ESS/hr associated with species tree leaf nodes of **N**_**e**_. Third, gene tree convergences were evaluated by their heights *h*_**G**_, their lengths *l*_**G**_, and the RNNI distances (Collienne and Gavryushkin, 2021) to their UPGMA *D*_*UPGMA*_ (Sokal, 1958) and neighbour-joining trees *D*_*NJ*_ (Saitou and Nei, 1987). As there are multiple gene trees, we only considered the mean ESS/hr of each term. Fourth, substitution model convergence (HKY substitution model; Hasegawa et al. (1985)) was measured from the transition-transversion ratio *κ*, nucleotide frequencies *f*, and gene tree substitution rates *ν*, where the ESS/hr of each term was averaged across all *k* substitution models. Lastly, relaxed clock model convergence was evaluated by considering the mixing of branch rate mean 𝔼 (**r**) and variance var(**r**), as well as the relaxed clock standard deviation parameter *σ*.

These results showed that, depending on the dataset, the “slowest” parameter generally converged considerably faster for StarBeast3 than it did for StarBeast2 (see the **min** term in Fig. S6–S10). On the smallest dataset considered (Frog), StarBeast2 and 3 performed comparably well overall (and no significant difference in **min**). However, StarBeast3 performed better on all of the other datasets, with the “slowest” parameter converging between 3.9 and 23 × as fast, and the posterior density *p*(*θ* | *D*) converging between 2.5 and 25 × as fast, usually at a statistically significant level.

Notably, relaxed clock model parameters converged up to 52 × as fast under StarBeast3. This was credited to the use of a real-space branch rate parameterisation (as opposed to the discrete branch rate categorisation employed by StarBeast2) as well as constant distance operators, which adjust branch rates and divergence times simultaneously (Zhang and Drummond, 2020; Douglas et al., 2021b). The disparity between StarBeast3 and StarBeast2 increased with the number of genes *k*, with one-and-a-half orders of magnitude improvement observed for the larger datasets, but only a 2-fold improvement for the *k* = 26 Frog data (Barrow et al., 2014), consistent with previous experiments (Douglas et al., 2021b).

Substitution model parameters *ψ* generally converged faster for StarBeast2 than they did for StarBeast3. Note, however, that this is by design. The total operator weight assigned to *ψ* parameters were 50% smaller in StarBeast3, in order to ensure balanced convergence across all areas of parameter space. In all datasets considered, substitution models converged significantly faster than any other area of parameter space, despite receiving relatively little operator weight, and therefore computational resources that were being spent on the substitution model were better off spent in “slower” areas of parameter space, such as gene tree node heights.

The `AdaptableOperatorSampler` operators (**Table S1**) confirmed the value in the `NER` and `ConstantDistanceMSC` operators for operating on their respective areas of parameter space. The `ConstantDistanceMSC` operator almost always outperformed other operators at proposing species node heights *t*_*S*_ (Table S4). The exception to this was the Skink dataset, for which the `UpDown` operator was superior at proposing branch lengths, and the Frog dataset, for which `ConstantDistanceMSC`, `CoordinatedExponential`, and `UpDown` were all on a par. In general, very little operator weight was rewarded to the `Uniform`, `Interval`, `TreeScale CoordinatedUniform`, and `CoordinatedExponential` operators for their abilities to propose species node heights. Similarly, among `NarrowExchange` variants evaluated by `AdaptableOperatorSampler (`*T*_*S*_), the `NER` operator was marginally favoured by all datasets (Table S5). This was due to the operator making larger or more frequent topological changes to the species tree, in faster computational runtime, especially compared with `CoordinatedNarrowExchange` and `CNER`. Overall, this experiment reinforced the value of learning operator weights on a problem-by-problem basis. A full breakdown of the remaining four adaptive operators can be found in Section 6 of *Supplementary Material*.

**Table S4:** Learned weights of the sub-operators of `AdaptableOperatorSampler(`*t*_*S*_), averaged across 5 replicates (2 sf). The operator which attained the highest proposal probability is indicated by a *.

| Dataset | Uniform | Interval | ConstantDistanceMSC | TreeScale | CUnif | CExp | UpDown |
| --- | --- | --- | --- | --- | --- | --- | --- |
| Frog | 0.06 | 0.0078 | 0.34* | 9.8e-05 | 0.04 | 0.22 | 0.33 |
| Simulated(12) | 1.1e-05 | 0.00013 | 0.99* | 2.6e-05 | 0.00043 | 8e-04 | 0.0061 |
| Simulated(48) | 3.9e-05 | 0.00049 | 0.98* | 5.8e-06 | 0.0011 | 3e-04 | 0.019 |
| Skink | 0.008 | 0.0087 | 0.34 | 4.8e-05 | 0.013 | 0.04 | 0.59* |
| Spider | 0.0019 | 0.0034 | 0.84* | 1.5e-05 | 0.0063 | 0.0025 | 0.15 |

**Table S5:**
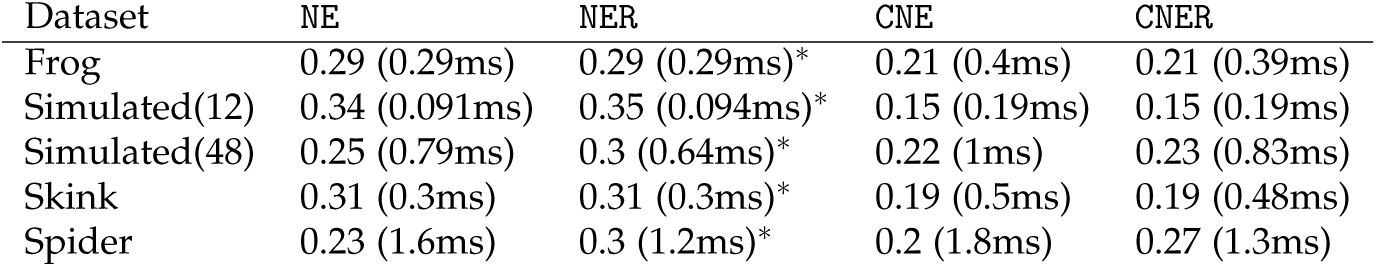
Learned proposal probabilities (and operator runtimes) for the suboperators of `AdaptableOperatorSampler` (*T*_*S*_), averaged across 5 replicates (2sf). Note that the timer starts at the beginning of the proposal and ends when the proposal has accepted or rejected. Notation: NE – narrow exchange; NER – narrow exchange rates; CNE – coordinated narrow exchange; CNER – coordinated narrow exchange rates. The operator which was rewarded the highest proposal probability for each dataset is indicated by a *.

Lastly, we evaluated the effect of threading on StarBeast3, by comparing its performance under 1, 2, 4, 8, and 16 threads allotted to the `ParallelMCMC` gene tree operator. There was a positive-but-modest correlation between the number of threads and the overall rate of convergence among the terms considered, with an overall log-linear slope coefficient of 0.19. This can be interpreted as follows: across the range of threads and datasets considered, doubling the number of threads was associated with an increase in mixing by 14%. Multithreading provided the strongest boost for the Skink and Spider datasets, and made little difference to the simulated dataset (48 taxa).

**Fig. S6:**
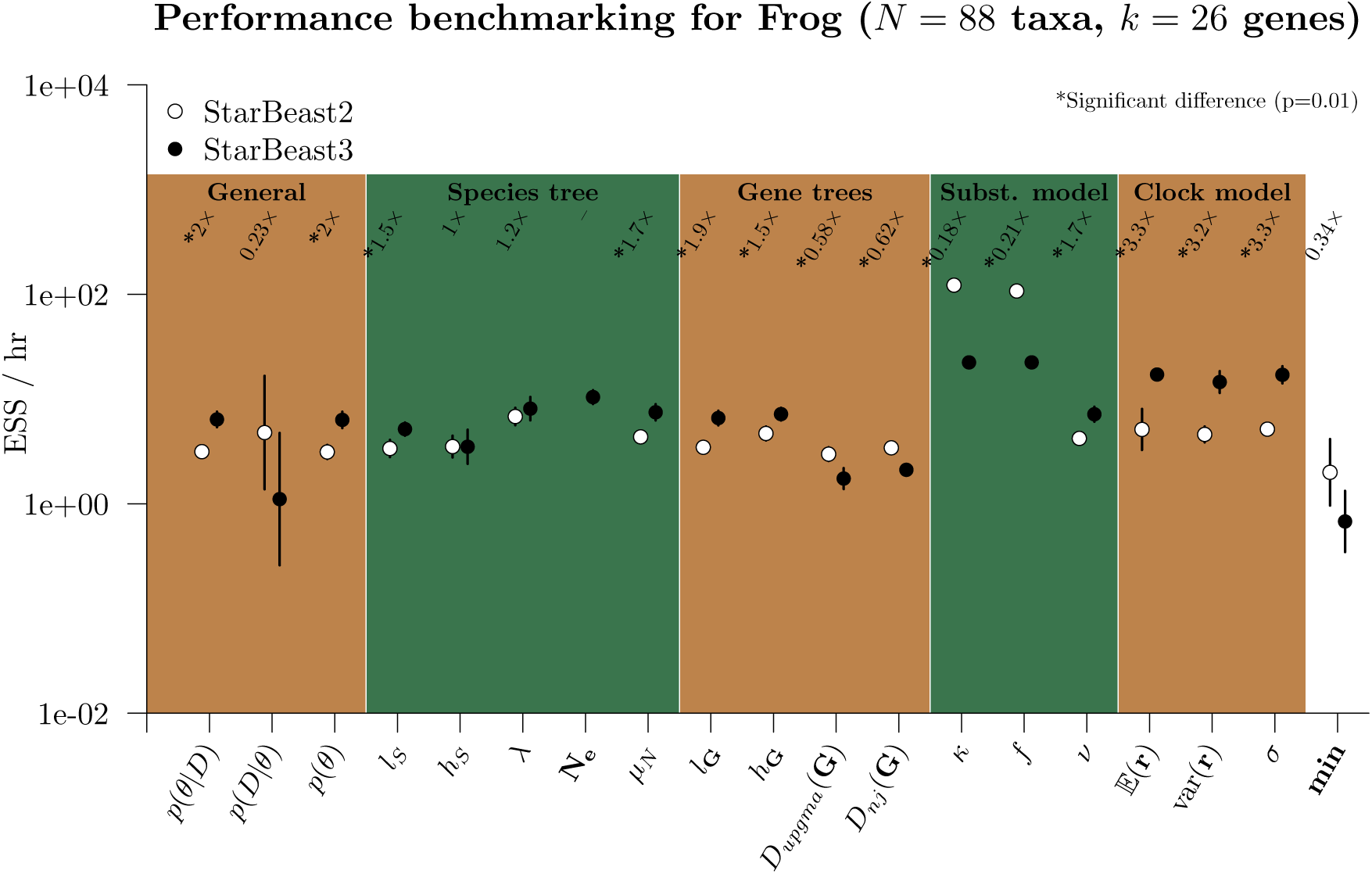
Performance benchmarking the Frog dataset (Barrow et al., 2014). Each point is the geometric-mean ESS/hr across 5 replicates, for either StarBeast2, or StarBeast3 with 16 threads. The geometric-mean relative performance of StarBeast3, compared with StarBeast2, is indicated above each term, and a * is present if the difference across 5 replicates is significant according to a Student t-test. Note that the y-axis is in log-space.

**Fig. S7:**
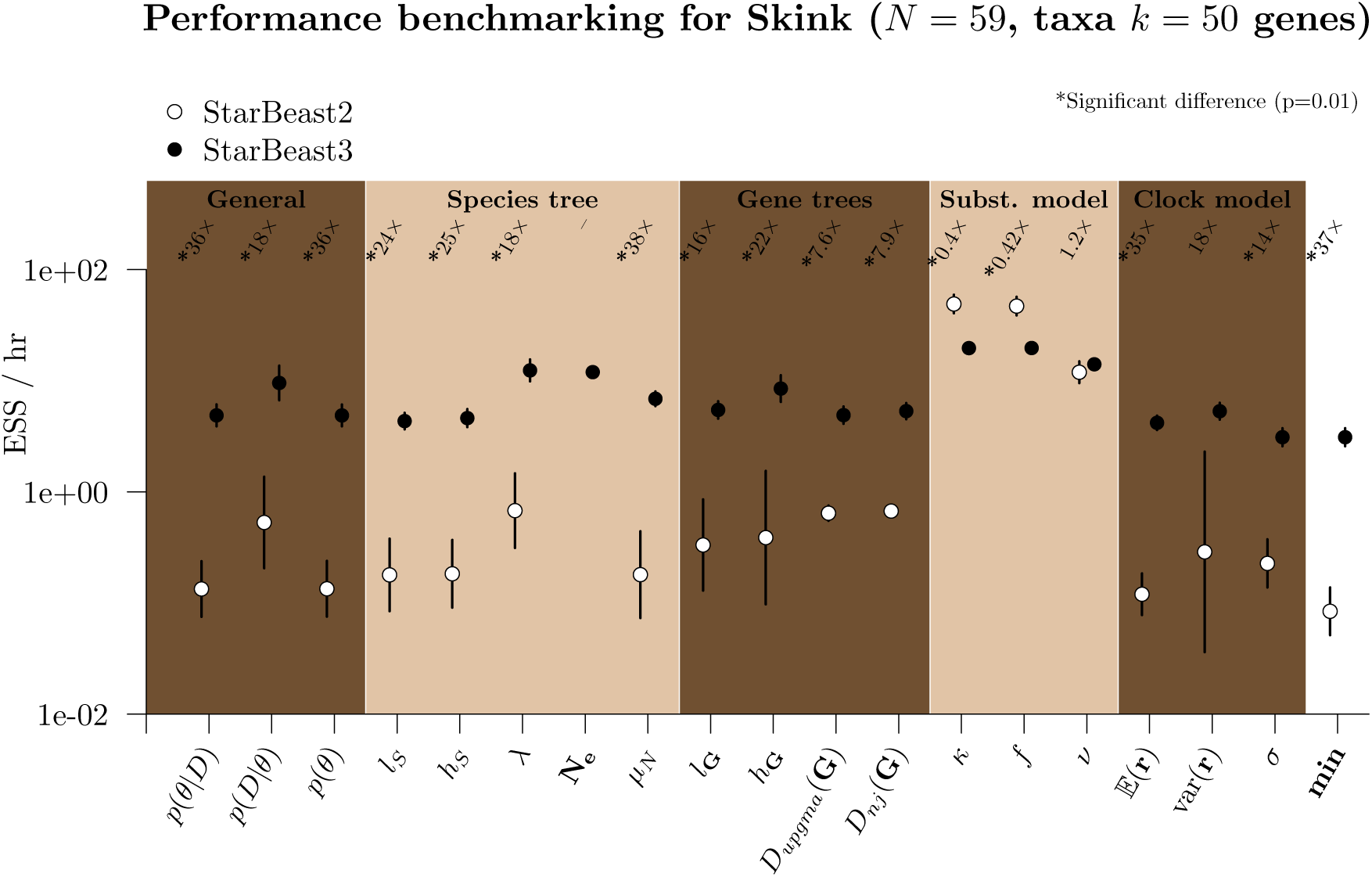
Performance benchmarking the Skink dataset (Bryson Jr et al., 2017). See Fig. S6 caption for figure notation.

**Fig. S8:**
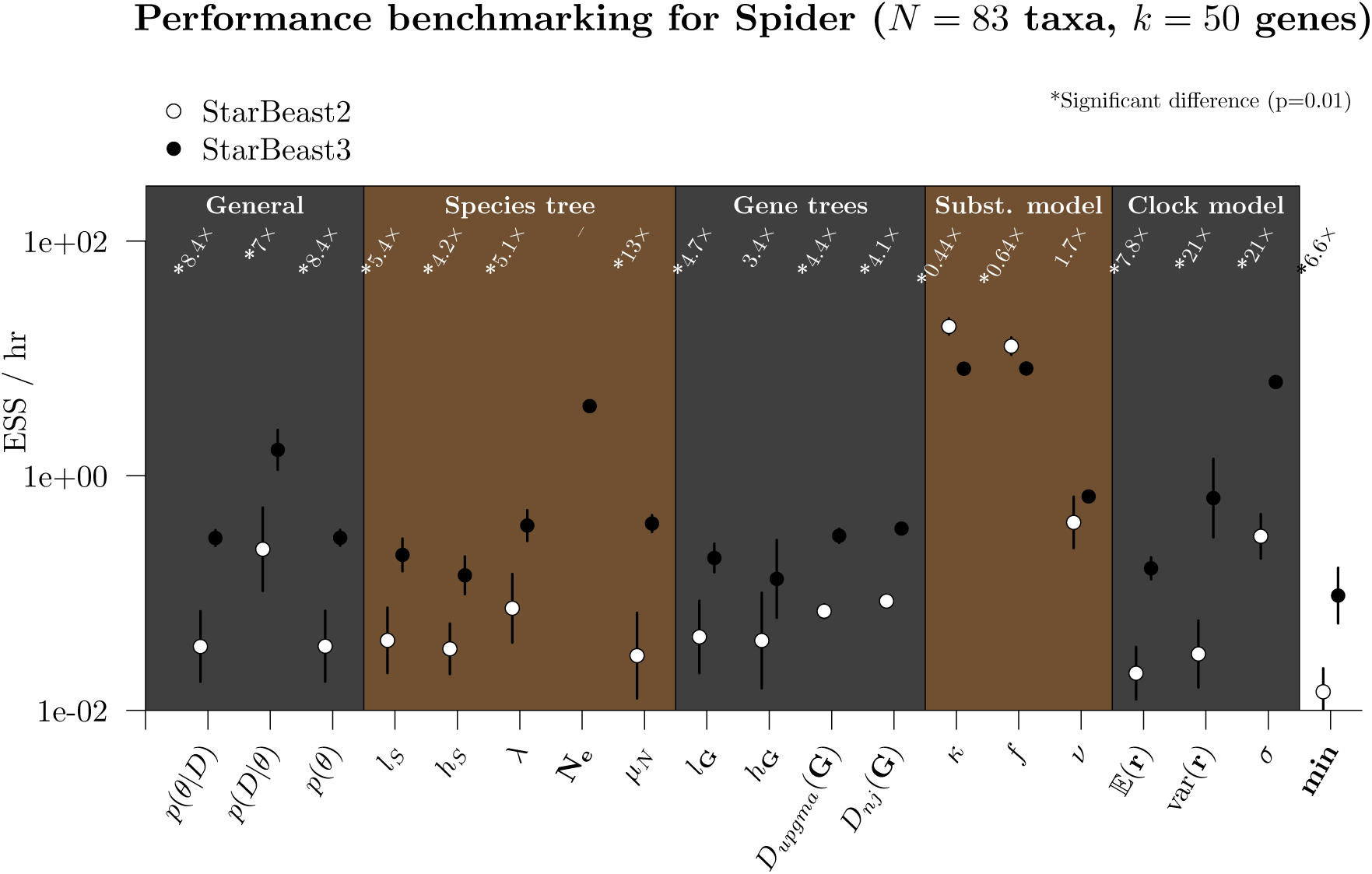
Performance benchmarking the Spider dataset (Hamilton et al., 2016). See Fig. S6 caption for figure notation.

**Fig. S9:**
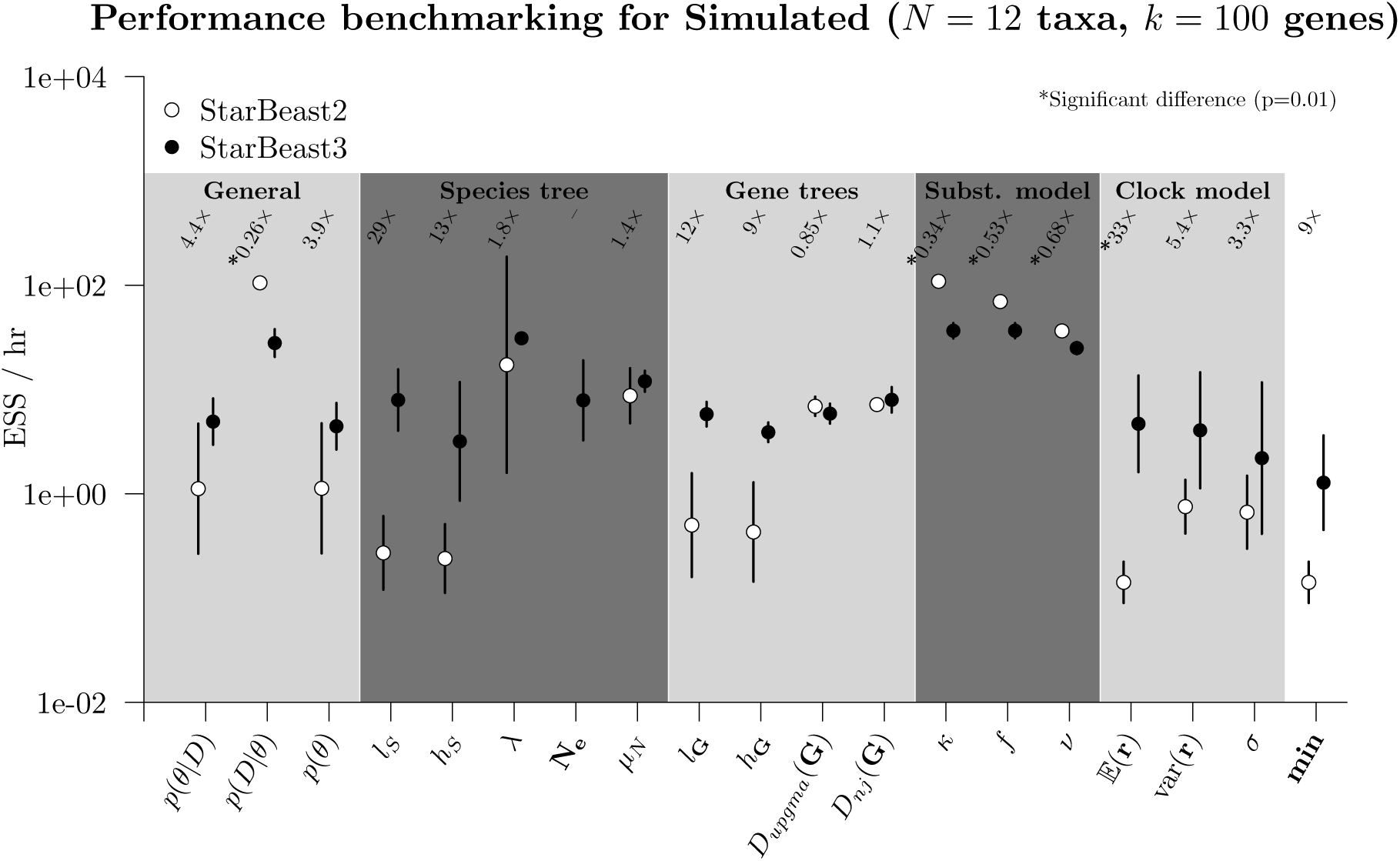
Performance benchmarking the 4 species Simulated dataset. See Fig. S6 caption for figure notation.

**Fig. S10:**
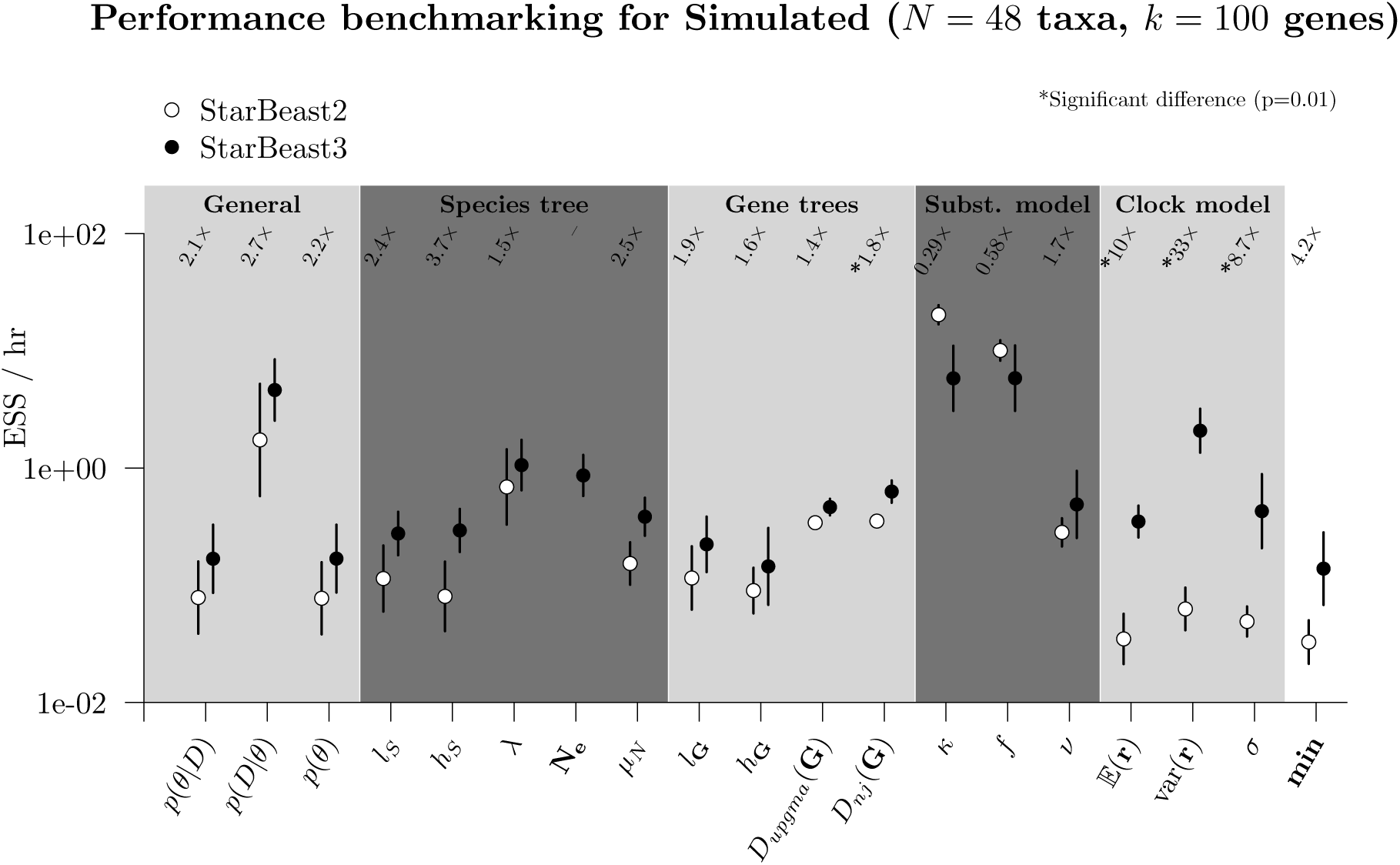
Performance benchmarking the 16 species Simulated dataset. See Fig. S6 caption for figure notation.

**Fig. S11:**
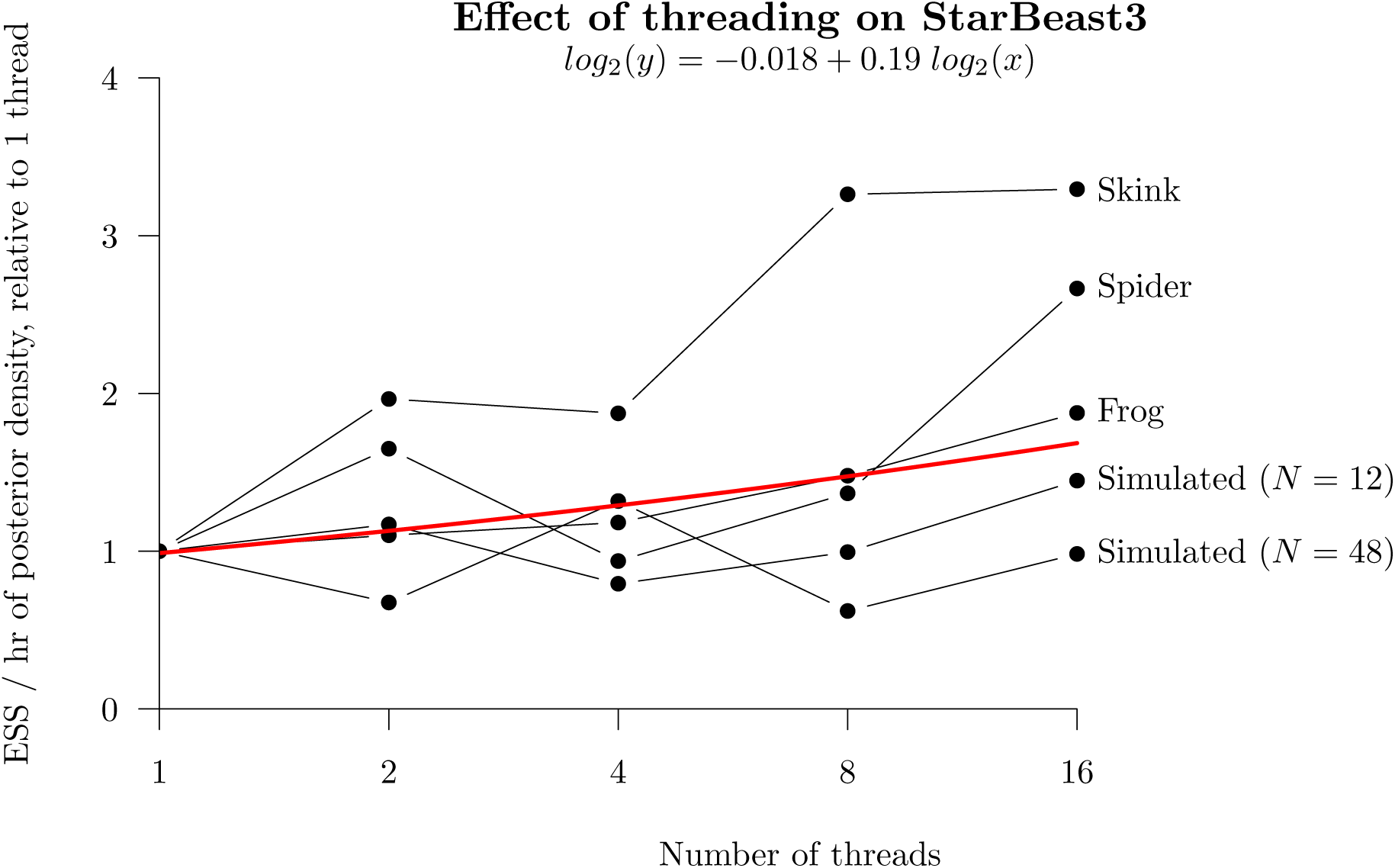
Effect of threading on StarBeast3 performance. Each point represents the ESS/hr of the posterior density *P*(*θ* | *D*) (averaged across 5 replicates), for the indicated thread count and dataset. These terms are normalised to enable comparison across datasets, by dividing it by that of 1 thread. A linear model was fit to the ESS/hr and number of threads, each in log_2_ space, and is reported at the top of the plot. The positive coefficient of the slope indicates that performance increased with the number of threads, across the range of threads considered. Parallel MCMC chain lengths were optimised using the adaptive scheme presented in Fig. S3.

## 4 Discussion

### 4.1 The Next Generation of Bayesian MCMC Operators

In recent years, Bayesian MCMC proposals have advanced significantly beyond that of the unidimensional random walk. The use of adaptive algorithms and advanced proposal kernels have become increasingly prevalent in recent years (Haario et al., 2001; Vihola, 2012; Benson et al., 2018; Yang and Rodríguez, 2013). In phylogenetic inference in particular, tree proposals have been guided by conditional clade probabilities and parsimony scores (Höhna and Drummond, 2012; Zhang et al., 2020), and mirror kernels learn target distributions which act as “mirror images” (Thawornwattana et al., 2018), for instance.

Here, we introduced a range of recently developed MCMC operators to the multispecies coalescent, including Bactrian proposal kernels (Yang and Rodríguez, 2013), which have been successfully applied to bird phylogeny (Maliet et al., 2019), and tree “flex” operators (BICEPS; Bouckaert (2021)), which have been applied to COVID-19 genomic data (Douglas et al., 2021a). We also invoked a series of more meticulous operators which account for known correlations, such as the AVMN kernel (Baele et al., 2017), constant distance operators (Zhang and Drummond, 2020), and the narrow exchange rate operator (Douglas et al., 2021b), as well as adaptive operators that improve over the course of MCMC, such as the adaptable operator sampler (Douglas et al., 2021b), parallel gene tree operators, and the AVMN kernel (Baele et al., 2017). Indeed, these operators have yielded a software package which outperforms StarBeast2 by up to one-and-a-half orders of magnitude, depending on the dataset and the parameter.

In order for Bayesian inference to keep up with the large volumes of genomic data, the development of efficient, meticulous, and adaptive MCMC operators is essential.

### 4.2 Efficient Parallelised Bayesian Inference of the Multispecies Coalescent

As genomic data becomes increasingly available, concatenating genomic sequences and inferring the phylogeny of the species as that of the genes can become enticing. However, this approach makes for an inconsistent estimator of topology when divergence times are small (Pamilo and Nei, 1988), and a biased estimator of species divergence times and substitution rates when incomplete lineage sorting is present (Arbogast et al., 2002; Ogilvie et al., 2016; Mendes and Hahn, 2016). Bayesian multispecies coalescent methods address these issues, but at the drawback of their demanding computational runtimes.

Therefore, as multithreading technologies become increasingly ubiquitous, the appeal in parallelising multispecies inference becomes clear. StarBeast3 exploits the assumption of conditional independence between gene trees, by doing Bayesian inference on gene trees in parallel, and therefore it scales with the size of the problem. StarBeast3 can process large datasets (100+ genes) and achieve convergence several times faster than its predecessors.

### 4.3 A Balanced Traversal Through Parameter Space

All areas of parameter space should be explored approximately evenly during MCMC. If one area of parameter space is being explored more rapidly than another, then computational resources allotted to the former should be diverted to the latter. This is best exemplified by the phylogenetic substitution model which, despite requiring relatively little attention to converge, still requires full recalculation of the tree likelihood upon every proposal (Felsenstein, 1981). Conversely, tree topologies often converge rather poorly and can require significant attention to be rescued from local optima. By fine tuning our MCMC operator proposal probabilities, we have achieved a balanced traversal through all areas of the multispecies coalescent parameter space. Although some parameters converge slower for StarBeast3 than they do for StarBeast2 (such as those in the substitution model), the slowest parameters converge significantly faster for the former; up to 30 × as fast (see the **min** term in Fig. S6 – S10).

For StarBeast3, we employed adaptable operators which are able to learn the proposal probabilities of other operators based on their ability to explore a single area of parameter space (Douglas et al., 2021b). However, there would be great benefit in an adaptable operator scheme which learns and applies a balanced exploration across different areas of parameter space on a problem-by-problem basis.

## 5 Conclusion

Here we introduce StarBeast3 – a software package for performing efficient Bayesian inference on genomic data under the multispecies coalescent model. We verified StarBeast3’s correctness and we benchmarked its performance against StarBeast2, which is an order of magnitude faster than its still popular predecessor *BEAST. We showed that StarBeast3 is significantly faster than StarBeast2. Notably, relaxed clock parameters converged up to one-and-a-half orders of magnitude faster, but most importantly even the “slowest” parameters converged up to 22 × faster. Our adaptive operator scheme allows proposal probabilities to be learned on a problem-by-problem basis, making StarBeast3 suitable for a range of datasets. By estimating effective population sizes (instead of analytically integrating the term out), we were able to parallelise gene tree proposals, and demonstrated that doubling the number of allotted threads was associated with an increase in performance by around 14%. StarBeast3 is highly effective at performing fast Bayesian inference on large datasets.

## 6 Software Availability

StarBeast3 is available as an open-source BEAST 2 package with an easy-to-use graphical user interface. Instructions for downloading and running StarBeast3 can be found at http://github.com/rbouckaert/starbeast3.

## 7 Acknowledgements

The study was supported by a Marsden grant 18-UOA-096 from the Royal Society of New Zealand. Software packages were benchmarked using the New Zealand eScience Infrastructure (NeSI) cluster, funded by the New Zealand Ministry of Business, Innovation and Employment.

