## Supplementary Information for "StarBeast3: Adaptive Parallelised Bayesian Inference of the Multispecies Coalescent"

#### SUPPLEMENTARY MATERIAL

### Contents

**1 Relaxed Clock Model Operators** **3**

**2 StarBeast2 Operator Scheme** **5**

**3 Prior Distributions** **6**

**4 Supplementary Well-Calibrated Simulation Study** **7**

**5 Evaluating the Performance Impact of New Operators** **9**
5.1 **GP**: Effective population size Gibbs sampler . . . . . . . . . . . . . . . . . . . . . . . . . . . . 9
5.2 **Real**: Relaxed clock model operators . . . . . . . . . . . . . . . . . . . . . . . . . . . . . . . . 9

**6 Supplementary Adaptive Operator Weighting Results** **10**

### List of Algorithms

S1 The `ConstantDistanceMSC` operator. . . . . 3
S2 The `CNER` operator. . . . . 4

### 1 Relaxed Clock Model Operators

The `ConstantDistanceMSC` operator can be found in **Algorithm S1** and the CNER operator can be found in **Algorithm S2**.

**Algorithm S1** The `ConstantDistanceMSC` operator. $t_i$ is the height (time) of node $i$, and $l_i$ is the length (time) of the branch above $i$. map$(x, i)$ returns the set of non-leaf nodes in gene tree $i$ which are contained within the branch above species node $x$.

**procedure** ConstantDistanceMsc($S$, $\mathbf{G}$, $\mathbf{N_e}$, $\mathbf{r}$)
  $\Sigma \leftarrow$ bactrian() ▷ Random walk follows Bactrian(0.95) distribution
  $X \leftarrow$ sampleNode($S$) ▷ Random non-leaf node
  $t_P \leftarrow t_X + l_X$ ▷ Parent height of $X$, or origin if tree if $X$ is root
  $(L, R) \leftarrow$ (leftChild($X$), rightChild($X$)) ▷ Children of $X$

  $t'_X \leftarrow t_X + \Sigma$ ▷ Propose new node height for $X$ using a Bactrian random walk

  ${r_X}' \leftarrow r_X \frac{t_P - t_X}{t_P - {t_X}'}$ ▷ Propose branch rates
  ${r_L}' \leftarrow r_L \frac{t_X - t_L}{t_X - {t_L}'}$
  ${r_R}' \leftarrow r_R \frac{t_X - t_R}{t_X - {t_R}'}$

  ${N_{e_X}}' \leftarrow N_{e_X} \frac{t_P - {t_X}'}{t_P - t_X}$ ▷ Propose population sizes
  ${N_{e_L}}' \leftarrow N_{e_L} \frac{t_P - {t_X}'}{t_P - t_X}$
  ${N_{e_R}}' \leftarrow N_{e_R} \frac{t_P - {t_X}'}{t_P - t_X}$

  **for** $i \in 1 \ldots k$ **do** ▷ Propose gene tree node heights
    **for** $\mathcal{X} \in \text{map}(X, i)$ **do** ▷ Gene nodes contained within $X$
      $t_{\mathcal{X}} \leftarrow t_P - (t_P - t_{\mathcal{X}}) \frac{t_P - t_X}{t_P - {t_X}'}$
    **for** $\mathcal{L} \in \text{map}(L, i)$ **do** ▷ Gene nodes contained within $L$
      $t_{\mathcal{L}} \leftarrow t_L + (t_{\mathcal{L}} - t_L) \frac{t_X - t_L}{t_X - {t_L}'}$
    **for** $\mathcal{R} \in \text{map}(R, i)$ **do** ▷ Gene nodes contained within $R$
      $t_{\mathcal{R}} \leftarrow t_R + (t_{\mathcal{R}} - t_R) \frac{t_X - t_R}{t_X - {t_R}'}$

**Algorithm S2** The `CNER` operator. Species tree nodes $A$, $B$, $C$, $D$, and $E$ have the following relationship: $E$ is the parent of $C$ and $D$, and $C$ is the parent of $A$ and $B$ in the original species tree before the proposal. `NarrowExchange`$([a_0, a_1], [b_0, b_1])$ proposes node $b_1$ as the parent of $a_0$ and node $a_1$ as the parent of $b_0$. See **Algorithm S1** for a description of map$(x, i)$.

**procedure** CNER$(A, B, C, D, E, \mathbf{G})$
    $A', B', C', D', E' \leftarrow$ `NarrowExchange`$([A, D], [C, E])$ ▷ Exchange $A$ with its uncle $C$

    $r'_A \leftarrow \frac{r_A(t_D - t_A) + r_D(t_E - t_D)}{t_E - t_A}$ ▷ Propose branch rates
    $r'_B \leftarrow \frac{r_B(t_D - t_B) + r_D(t'_D - t_D)}{t'_D - t_B}$
    $r'_C \leftarrow \frac{r_C(t_E - t_C) - r_D(t_E - t'_D)}{t'_D - t_C}$

    **for** $i \in 1 \ldots k$ **do** ▷ Regraft gene trees
        **for** $\mathcal{D} \in$ map$(D, i)$ **do** ▷ Get all nodes in $g_i$ that map to $D$

            ▷ Regraft $\mathcal{D}$ onto a branch in $D'$ if exactly one child subtree maps to $A$
            AContainsLeft $\leftarrow \exists\, j \in$ subtree(leftChild$(\mathcal{D})$) : $j \in$ map$(A, i)$
            AContainsRight $\leftarrow \exists\, j \in$ subtree(rightChild$(\mathcal{D})$) : $j \in$ map$(A, i)$

            **if** AContainsLeft $\oplus$ AContainsRight **then**
                $X \leftarrow$ sample(map$(D', i)$)
                ${g_i}' \leftarrow$ `NarrowExchange`$([\mathcal{D}, \text{parent}(\mathcal{D})], [X, \text{parent}(X)])$ ▷ Regraft $\mathcal{D}$

### 2 StarBeast2 Operator Scheme

The operator scheme used by StarBeast2 during all benchmarks studied here is presented in Table S1. This is the default StarBeast2 operator scheme as generated by BEAUti. We used StarBeast2 v0.15.13 throughout this study.

| Operator | Weight | Reference |
|---|---|---|
| **General / species tree** | 20% | |
| Scale($\mu_N$) | 1 | |
| Scale($\lambda$) | 1 | |
| NodeReheight($T_S, t_S, T_{\mathbf{G}}, t_{\mathbf{G}}$) | 30 | Ogilvie et al. (2017) |
| CoordinatedUniform($t_S, t_{\mathbf{G}}$) | 15 | Ogilvie et al. (2017); Jones (2017) |
| CoordinatedExponential($t_S, t_{\mathbf{G}}$) | 15 | Ogilvie et al. (2017); Jones (2017) |
| SubtreeSlide($T_S, t_S$) | 15 | Hohna et al. (2008) |
| WilsonBalding($T_S$) | 15 | Drummond et al. (2002) |
| WideExchange($T_S$) | 15 | Drummond et al. (2002) |
| NarrowExchange($T_S$) | 15 | Drummond et al. (2002) |
| Uniform($t_S$) | 15 | |
| RootScale($t_S$) | 3 | |
| Scale($t_S$) | 3 | |
| UpDown([$t_S, t_{\mathbf{G}}, \mu_N$], [$\nu, \lambda, \mu$]) | 6 | Drummond et al. (2002) |
| **Gene trees / site models / clock model** | 80% | |
| $\forall_{i \in 1,\ldots,k}$ SubtreeSlide($T_{g_i}, t_{g_i}$) | 15 | Hohna et al. (2008) |
| $\forall_{i \in 1,\ldots,k}$ WilsonBalding($T_{g_i}$) | 15 | Drummond et al. (2002) |
| $\forall_{i \in 1,\ldots,k}$ WideExchange($T_{g_i}$) | 15 | Drummond et al. (2002) |
| $\forall_{i \in 1,\ldots,k}$ NarrowExchange($T_{g_i}$) | 15 | Drummond et al. (2002) |
| $\forall_{i \in 1,\ldots,k}$ Uniform($t_{g_i}$) | 15 | |
| $\forall_{i \in 1,\ldots,k}$ RootScale($t_{g_i}$) | 3 | |
| $\forall_{i \in 1,\ldots,k}$ Scale($t_{g_i}$) | 3 | |
| $\forall_{i \in 1,\ldots,k}$ UpDown([$t_{g_i}$], []) | 3 | Drummond et al. (2002) |
| $\forall_{i \in 1,\ldots,k}$ Scale($\kappa_i$) | 3 | |
| $\forall_{i \in 1,\ldots,k}$ Scale($\nu_i$) | 1 | |
| $\forall_{i \in 1,\ldots,k}$ DeltaExchange($f_i$) | 1.5 | |
| Scale($\sigma$) | 1 | |
| Uniform($\mathbf{r}$) | 9 | |
| Cycle($\mathbf{r}$) | 9 | Ogilvie et al. (2017) |

**Table S1:** StarBeast2 operator scheme, assuming a Yule tree prior on the species tree with birth rate $\lambda$. Operator weights are renormalised such that those under the **General / Species tree** scheme have a cumulative weight of 20%, while the remaining operator weights are a function of $k$ and sum to 80%.

### 3 Prior Distributions

The prior distributions used in all validation and benchmarking analyses throughout this article are summarised in Table S2. The ploidy of each population is assumed to be 2 throughout this article.

| Parameter | Description | Prior Distribution |
|---|---|---|
| $\lambda$ | Yule-model birth rate | LogNormal($\mu = 1, \sigma = 1.25$) |
| $\kappa$ | Transition-transversion ratio | LogNormal($\mu = 1, \sigma = 1.25$) |
| $f$ | Nucleotide frequencies | Dirichlet($\alpha = (10, 10, 10, 10)$) |
| $\nu$ | Gene tree rate | LogNormal($\mu = -0.18, \sigma = 0.6$)$^{\dagger}$ |
| $\sigma$ | Relaxed clock standard deviation | Gamma($\alpha = 0.5396, \beta = 0.3819$) |
| $\mathbf{r}$ | Branch rates | LogNormal($\mu = -\frac{\sigma^2}{2}, \sigma = \sigma$)$^{\dagger}$ |
| $\mu_N$ | Effective population size mean | LogNormal($\mu = -5, \sigma = 1.25$) |
| $\mathbf{N_e}$ | Effective population size | InverseGamma($\alpha = 2, \beta = \mu_N$) |

**Table S2:** Prior distributions. A LogNormal($\mu, \sigma$) distribution has a mean of $\mu$ and a standard deviation of $\sigma$ in log-space. $^{\dagger}$ $\mu$ and $\sigma$ are set such that the distribution has a mean of 1 in real-space.

### 4 Supplementary Well-Calibrated Simulation Study

We performed an additional well-calibrated simulation studies to validate StarBeast3. This was achieved by considering $n_S = 4$ species, $n_{\mathbf{G}} = 12$ taxa, and $k = 100$ genes (Fig. S1).
The 95%-coverage of each parameter was approximately 95% (meaning that the true parameter estimate was within the 95% highest posterior density interval approximately 95% of the time). In addition to the validation presented in the main article, this provides confidence that StarBeast3 is correctly implemented and is able to recover parameter estimates during Bayesian MCMC. Each well-calibrated simulation here and in the main article was run until the effective sample size of each reported parameter was over 200, demonstrating convergence.

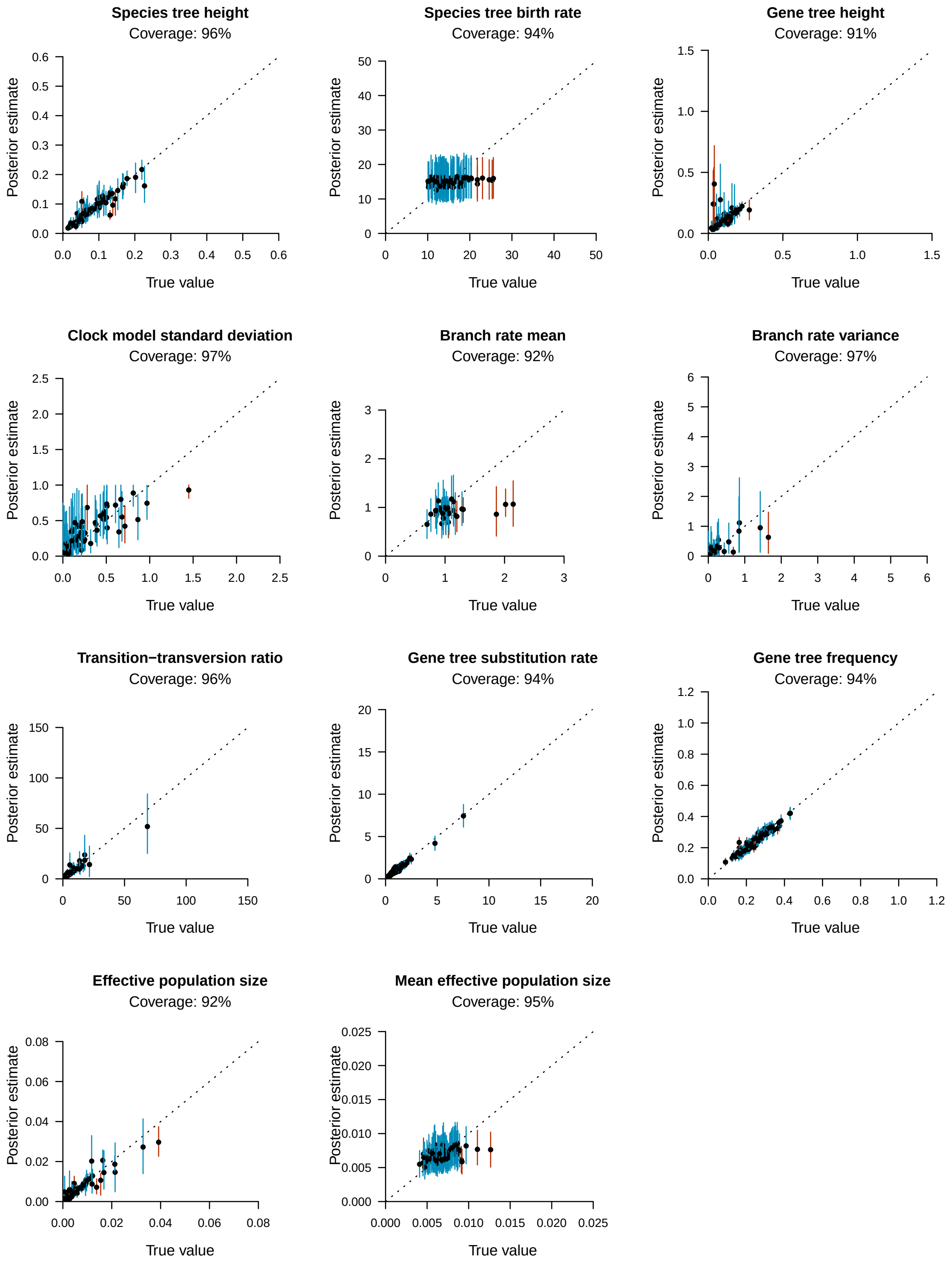

**Fig. S1:** Well-calibrated simulation study for $n_S = 4$ species, $n_G = 12$ taxa, and $k = 100$ genes. 100 simulations were performed to recover the coverage between "true" simulated values and their estimates under the posterior distribution. 95% highest posterior density (HPD) intervals of parameters are represented by vertical lines. Each line represents a single simulation, and is coloured blue when the true value was contained within the 95% interval, or red otherwise. The top of each plot shows the coverage of each parameter (i.e., the number of MCMC simulations for which the "true" parameter value was contained within the 95% HPD).

### 5 Evaluating the Performance Impact of New Operators

The AVMN and Bactrian kernels have previously been benchmarked, and were found to be up to an order of magnitude faster (Baele et al., 2017), and 10-50% faster than standard random walks, respectively (Yang and Rodríguez, 2013; Douglas et al., 2021). Here, we incrementally introduce newly developed operators onto the default StarBeast2 settings in order to evaluate the impact that each operator has on parameter convergence during Bayesian MCMC. This was measured by computing effective sample size (ESS) generated per hour during MCMC across 5 replicates of the $N = 12$ taxa Simulated dataset.

#### 5.1 GP: Effective population size Gibbs sampler

We evaluated the performance of `GibbsPopulation` operator at estimating population sizes, compared with the standard `Swap` and `Scale` operators employed by StarBeast2. The former operator scheme is referred to as **GP** and the latter is **Scale**. All other operators, including their proposal probabilities, remained constant for both settings, and a strict clock model was employed (i.e., $\mathbf{r} = \mathbf{1}$). These results showed that the ESS/hr of $\mathbf{N_e}$ (leaves only) was an average of $1.1\times$ times as large, and that of the posterior density $p(\theta|D)$ was $1.1\times$ as large for the former setting than the latter, thus confirming the effectiveness of this new operator. Because this operator acts on all population sizes simultaneously, the performance improvement is likely to be even greater for larger species trees.

#### 5.2 Real: Relaxed clock model operators

We introduced the relaxed clock model and its new operators (`AdaptableOperatorSampler(`$\mathbf{r}$`)` and `AdaptableOperatorSampler(`$\sigma$`)`) to **GP**, and compared it with the standard StarBeast2 relaxed clock model operator scheme (building on top of **Scale**). The former branch rates were parameterised as real numbers (**Real**) while the latter were modelled as discrete categories (**Cat**). All other operators, including their proposal probabilities, remained constant. These results showed that the ESS/hr of **Real+GP** were an average of $10\times$ as large for $\mathbb{E}(\mathbf{r})$, $14\times$ for $\text{var}(\mathbf{r})$, $17\times$ for $\sigma$, $5.2\times$ for $l_S$, $2.4\times$ for the posterior density $p(\theta|D)$, compared with **Cat+Scale**. Overall, the real-space relaxed relaxed clock parameterisation and its operators were a significant improvement.

### 6 Supplementary Adaptive Operator Weighting Results

The operator weights learned by each `AdaptableOperatorSampler` are presented in this section. As expected, the optimal operator for any given parameter is dependent on the dataset, however most benchmarked datasets were in general agreement regarding which operator performs the best.

Unidimensional parameters (such as species tree birth rate $\lambda$, population size mean $\mu_N$, and the relaxed clock standard deviation $\sigma$) were most efficiently operated on by their respective `Scale` operators (Tables S3, S4, S5). However, the simple `SampleFromPrior` operator was often successful at resampling parameters from their prior, with proposal acceptance rates reaching $> 20\%$ on some datasets and $< 1\%$ on others.

It is important to emphasise that the operators favoured among the datasets considered are unlikely to be the same among future datasets.

| Dataset | `Scale` | `UpDown` | `SampleFromPrior` |
|---|---|---|---|
| Frog | 0.99* | 8.8e-06 | 0.01 (1.2%) |
| Simulated(12) | 0.88* | 1.4e-06 | 0.12 (23%) |
| Simulated(48) | 0.95* | 3.2e-07 | 0.049 (4.2%) |
| Skink | 1* | 5.5e-06 | 0.00046 (0.1%) |
| Spider | 1* | 3.3e-06 | 0.00063 (0.086%) |

**Table S3:** Learned weights of the sub-operators of `AdaptableOperatorSampler(`$\lambda$`)`, averaged across 5 replicates (2 sf). The operator which attained the highest proposal probability is indicated by a *. The average proposal acceptance percentage is indicated for `SampleFromPrior`.

| Dataset | `Scale` | `UpDown` | `SampleFromPrior` |
|---|---|---|---|
| Frog | 0.82* | 0.0037 | 0.18 (8.9%) |
| Simulated(12) | 0.64* | 0.00045 | 0.36 (25%) |
| Simulated(48) | 0.77* | 0.00029 | 0.23 (13%) |
| Skink | 0.83* | 0.003 | 0.16 (7.8%) |
| Spider | 0.84* | 0.0042 | 0.15 (7.5%) |

**Table S4:** Learned weights of the sub-operators of `AdaptableOperatorSampler(`$\mu_N$`)`, averaged across 5 replicates (2 sf). The operator which attained the highest proposal probability is indicated by a *. The average proposal acceptance percentage is indicated for `SampleFromPrior`.

| Dataset | `Scale` | `SampleFromPrior` |
|---|---|---|
| Frog | 0.85* | 0.15 (8.7%) |
| Simulated(12) | 0.91* | 0.089 (4.8%) |
| Simulated(48) | 0.86* | 0.14 (7.4%) |
| Skink | 0.81* | 0.19 (12%) |
| Spider | 0.98* | 0.021 (0.88%) |

**Table S5:** Learned weights of the sub-operators of `AdaptableOperatorSampler(`$\sigma$`)`, averaged across 5 replicates (2 sf). The operator which attained the highest proposal probability is indicated by a *. The average proposal acceptance percentage is indicated for `SampleFromPrior`.

| Dataset | `Scale` | `ConstantDistanceMSC` | `SampleFromPrior` |
|---|---|---|---|
| Frog | 0.29 | 0.47* | 0.24 (23%) |
| Simulated(12) | 0.015 | 0.94* | 0.05 (8.2%) |
| Simulated(48) | 0.03 | 0.86* | 0.11 (15%) |
| Skink | 0.28 | 0.46* | 0.26 (26%) |
| Spider | 0.032 | 0.93* | 0.041 (20%) |

**Table S6:** Learned weights of the sub-operators of `AdaptableOperatorSampler(r)`, averaged across 5 replicates (2 sf). The operator which attained the highest proposal probability is indicated by a *. The average proposal acceptance percentage is indicated for `SampleFromPrior`.
